# Vision-Based Collective Motion: A Locust-Inspired Reductionist Model

**DOI:** 10.1101/2023.01.17.524210

**Authors:** David L. Krongauz, Amir Ayali, Gal A. Kaminka

## Abstract

Naturally occurring collective motion is a fascinating phenomenon in which swarming individuals aggregate and coordinate their motion. Many theoretical models of swarming assume idealized, perfect perceptual capabilities, and ignore the underlying perception processes, particularly for agents relying on visual perception. Specifically, biological vision in many swarming animals, such as locusts, utilizes monocular non-stereoscopic vision, which prevents perfect acquisition of distances and velocities. Moreover, swarming peers can visually occlude each other, further introducing estimation errors. In this study, we explore necessary conditions for the emergence of ordered collective motion under restricted conditions, using non-stereoscopic, monocular vision. We present a model of vision-based of collective motion for locust-like agents: elongated shape, omni-directional visual sensor parallel to the horizontal plane, and lacking stereoscopic depth perception. The model addresses (i) the non-stereoscopic estimation of distance and velocity, (ii) the presence of occlusions in the visual field. We consider and compare three strategies that an agent may use to interpret partially-occluded visual information at the cost of the computational complexity required for the visual perception processes. Computer-simulated experiments conducted in various geometrical environments (toroidal, corridor, and ring-shaped arenas) demonstrate that the models can result in an ordered or near-ordered state. At the same time, they differ in the rate at which order is achieved. Moreover, the results are sensitive to the elongation of the agents. Experiments in geometrically constrained environments reveal differences between the models and elucidate possible tradeoffs in using them to control swarming agents. These suggest avenues for further study in biology and robotics.

**Author summary:** Swarm collective motion is a wide-ranging phenomenon in nature, with applications in multi-agent, multi-robot systems. In most natural swarming species, individuals rely on monocular, non-stereoscopic vision as the key sensory modality for their interactions. For example, the migratory locust (*locusta migratoria*) displays large swarms of individuals, moving in alignment and relying solely on non-stereoscopic visual perception. Inspired by these locust swarms, we have developed a monocular, non-stereoscopic vision-based model that achieves synchronized motion in a swarm of two-dimensional agents, even with inaccurate estimates of distances and velocities, particularly in the presence of occlusions. We explore three general strategies for handling occlusions, which differ in the requirements they place on the complexity of the visual perception process. We show that strategies may reach a highly ordered motion state but differ in their convergence rate.

## 1 Introduction

Swarms composed of large groups of individuals can engage in coordinated collective motion, without centralized or group-wide control, global perception, or global communications. This coordinated collective motion (which we henceforth term *flocking*, but is also known as *schooling*, or *swarming*) describes the emergence of a common heading for the motion of agents in the swarm. Flocking can arise from disordered initial conditions, where initial headings and positions are arbitrary, despite the restricted locality of the perception and action of any individual agent in the swarm.

In nature, flocking is ubiquitous in birds [1, 2], fish [3, 4], insects [5, 6], bacteria [7], and human crowds [8–12]. It is a phenomenon that has been of interest to the scientific community for decades, inspiring modeling efforts (e.g., [13, 14]) and bio-mimetic technologies in graphics [13, 15]), simulations (e.g., [16, 17]), and robotics (see [18] for a recent survey).

The leading paradigm underlying models of collective motion is that it results from repeated local (myopic) interactions among individual swarm members (see, e.g., [19, 20]). The control procedure of each single agent translates its perception of the local physical and social (nearby conspecifics) environments into a decision regarding its next action. The local, individual decisions of numerous agents—interacting with each other—result in the eventual convergence of the group to an ordered state, in which the agents head in a common heading (that may change dynamically). Commonly, flocking agents are modeled as *self-propelled particles* (SPP) that are continuously subjected to the mutual steering forces caused by their neighbors [21]. These mutual force interactions feed into the agents’ decision-making, changing their motion [14, 22, 23]. Under appropriate conditions, this generates flocking [24–26].

Traditional models of flocking abstract away from the real limitations of perceptual processes. They rely on idealized perceptual capabilities that allow agents to determine their neighbors’ distances, headings, and velocities (see, for instance, [13, 14, 19, 20, 27]). This ignores the sensory and computational limitations inherent to physical agents in nature or in a robotics laboratory: limited effective sensing regions (width and range of the sensory field of view), systematic perceptual ambiguities, computational resources required for sensor information processing, and sensitivity to occlusions of some neighbors by others (common in flocking) [28].

In living species or in robots that use vision as a primary sensory modality, the underlying sensory structure and processing abilities of the agent places multi-faceted constraints on the possible visual perception processes that may be employed. The position of the eyes/visual sensors, and the angular and range limitations on their fields of view, constrain the perception strategies that can be used to provide the information needed for flocking. These strategies vary in accuracy, failure modes, and computational/cognitive complexity they demand of the individual brain [29–31].

For example, when an agent has two or more sensors that have intersecting fields of view, *stereopsis* (stereoscopic vision) can be used to accurately estimate distance—but the intersecting field of view is relatively narrow, and its effective range is short [32]. In contrast, when one or more eyes generate monocular (non-stereoscopic) images, distances may be inferred by matching conspecific visual templates, by integrating images over time to compute optical flow, or by other strategies [33–40], all of which vary in computational requirements and accuracy. The tradeoffs involved, their biological plausibility, their potential computational costs, and the opportunities they offer for robots are currently not well understood.

Marching locust nymphs [41–43] offer an inspiring example to challenge our understanding of vision-based collective motion. The individual locust nymph lacks binocular depth perception, though its two eyes offer an almost-perfect omni-directional visual field. Both field and laboratory studies indicate that the robust locust collective motion emerges from the interactions between individuals [26, 44–46]. It is largely accepted that non-stereoscopic vision is the key sensory modality underlying these local interactions. With limited processing power, and having no depth perception, the individual locust makes motion decisions based on visual information that lacks precision in measurement of its neighbors’ proximity, headings, or velocities. Despite these limitations, locusts display impressive flocking, involving large numbers of individual agents. Models that ignore the visual perception processes lack the explanatory power to capture how this is achieved.

Recent studies of monocular vision-based flocking have investigated some relevant related mechanisms. Studies of natural swarms (often in vertebrates) [3, 28, 47–50] and robot swarms [51–53] have suggested strategies for forming dynamic *sensory* networks, by which agents remain connected to each other while attending to only a subset of their neighbors at any given time. These are useful both in cases of a limited field of view, and in handling the occlusions that limit the ability to recognize and track neighbors that are only partly visible. Other studies have focused on the mechanisms used by the individual for visual processing, given a specific morphology of agents [54–59]. The different studies all reveal important insights but often make assumptions (e.g., that agents are circular, or that they can sense the orientation of visible neighbors), that may not be relevant for the locust body morphology or of its perception capabilities (we discuss these investigations in more depth in Section 5).

Inspired and challenged by the marching locust phenomenon, we have developed a *reductionist* model of monocular, non-stereoscopic, vision-based collective motion in *locust-like* agents (Section 2). The model builds on the geometrical characteristics of locust body morphology and visual perception (wide field of view, monocular images, elongated shape), but reduces the visual inputs to the bare minimum perceivable in two dimensions (i.e., no height information is used; objects are perceived only along the horizontal sensory plane). We present a control algorithm that employs only the information accessible via the agent’s visual field (Section 2.1). We then propose several general strategies that the agent might employ when assessing partially obstructed neighbors (Section 2.2). From these restricted capabilities, the control algorithm synthesizes flocking under various environment conditions and occlusion-handling strategies.

Experiments performed via computer simulation (Section 3) explored the emergence of ordered (flocking) movement under various conditions: varying group sizes, range of the visual field, body lengths, and strategies for handling occlusions. The experiments were performed in various simulated arenas, that differed in their border periodicity constraints and area.

Our goal was to elucidate strategies that organisms—and robot builders—can use to trade computation or cognitive complexity for reliable ordered collective motion. The results (Section 4) reveal that in many cases, the swarm’s order parameter, which characterizes the level of alignment between the agents in the swarm, reaches high values regardless of occlusion-handling strategy or environment constraints. However, in highly constrained arenas, different strategies for handling occlusions differ in the rate and degree of emerging order. Furthermore, the body morphology (specifically, body elongation) impacts the rate in which order is achieved, in different strategies. Section 5 presents an in-depth discussion of the results, their relation to previous models, and their implications for future research efforts.

## 2 A Reductionist, Non-Stereoscopic Model of Visual Perception for Collective Motion

We present a reductionist model of non-stereoscopic vision-based collective motion, from the perspective of a locust-like agent. First, in Section 2.1, we present the restricted visual perception mechanisms, and the vision-based algorithm governing the agents’ movement. Next, in Section 2.2, we discuss the potentially harmful effects of occlusions on perception. We then present three alternative strategies allowing the algorithm to interpret and deal with partially occluded visual information.

### 2.1 The Principal Monocular Vision-Based Flocking Model

We begin with the basic geometry of the agent. We consider a group of *N* identical rectangular agents with width *w* and length *l*, moving in a two-dimensional environment at velocity ***v***_***i***_, parallel to their length axis. The elongation of the agents is measured by the ratio of length to width (*l/w*), such that the ratio is ≥1, i.e., a square is the shortest agent.

The position coordinates ***x***_***i***_ of agent *i* are updated at discrete time steps according to the motion equation,

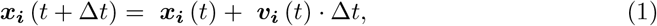

with velocity ***v***_***i***_(*t*) updated at each time step causing the agent to steer towards a desired velocity with steering-parameter factor *η*,

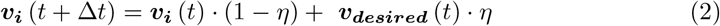

***v***_***desired***_ is calculated based on the decision algorithm of the Vicsek Model [14].

Assuming agent *i* has a set of neighbors *J*_*i*_, its desired velocity averages the velocities of the neighbors *j ∈ J*_*i*_ at each time *t*:

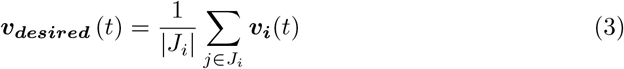

The question, of course, is how the velocities of neighbors are estimated based on visual information. To explore this in-depth, we first discuss the geometry of locust-like visual perception.

#### The Geometry of Locust-Like Vision

We model each agent’s visual field of view by an idealized omnidirectional sensor covering 360 degrees around the observing agent (hereafter, the *focal agent*). This wide field of view is consistent with the nearly omni-directional field of view of locust nymphs [60, 61]. The range of the sensing, i.e., the maximum distance at which it can detect other agents, is denoted *R*, a parameter of the model (which we examine in the experiments).

Figure 1a presents the basic geometry and notation for a focal agent *o* heading “up”, with velocity vector ***v***_***o***_. The focal agent has a single neighbor *j* moving with velocity ***v***_***j***_ and located at a distance *r*_*j*_ *< R* measured between the focal agent and the neighbor *j* along the line of sight (LOS) connecting the center-of-mass (COM) of the neighbor (*COM*_*j*_) and the COM of the focal agent (*COM*_*o*_). We denote the displacement *vector* of neighbor *j* equals ***r***_***j***_ = *COM*_*j*_ *− COM*_*o*_, while the *scalar* distance to *j* is *r*_*j*_ = *∥****r***_***j***_*∥*. The velocity ***v***_***j***_ is composed of the tangential velocity ***v***_***j***,***t***_ and radial velocity ***v***_***j***,***r***_ components, relating to the line of sight (LOS). The angular position of the neighbor *j* relative to the heading direction is denoted as (bearing; *β*_*j*_), and the angle subtended on *o*’s visual sensor is denoted as *α*_*j*_. This angle is calculated as an angle between the *edge rays* from the focal agent to the *observed* corners of the neighbor [62]. The edge rays mark the maximally-distant observable pair of the neighbor’s corners. The line segment connecting these two corners is called the *effective edge*. Fig. 1b illustrates the edge rays for three neighbors observed by the focal agent.

**Fig 1.**
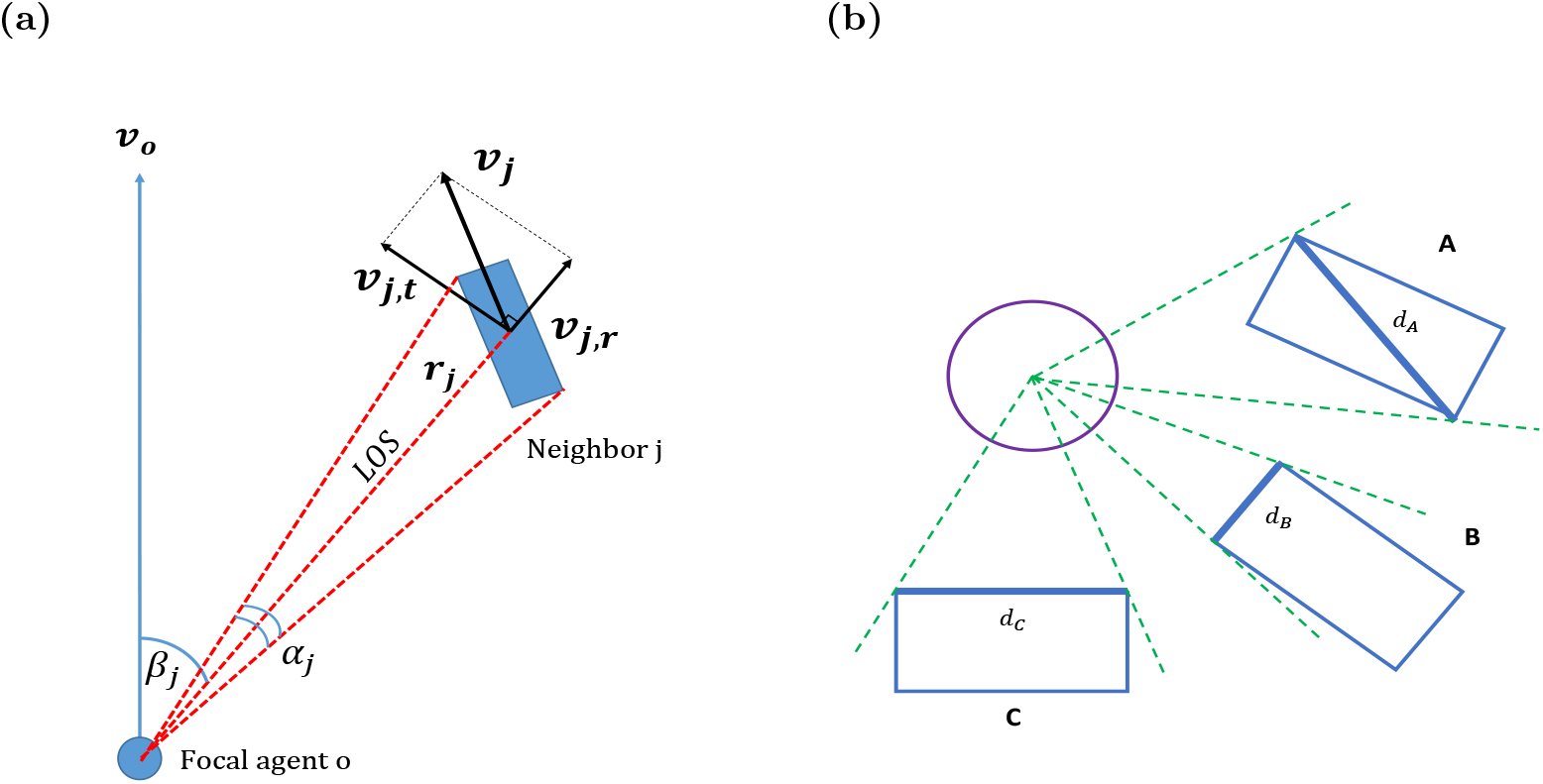
Notations and Definitions. **(a)** A schematic depiction of a neighbor’s observed geometrical features and notation used. The bearing angle *β*_*j*_ defines the angle between the heading of the focal agent (*v*_*o*_) and the line of sight (LOS, as defined in the text). The circle represents the idealized sensor of the focal agent. The subtended angle *α*_*j*_ is defined as the angle between the edge rays directed towards the extremities of the neighbor. The distance from the focal agent to the center of neighbor j is denoted *r*_*j*_. The neighbor’s velocity ***v***_***j***_, is composed of two orthogonal components: the radial component ***v***_***r***,***j***_ is parallel to LOS, and the tangential component ***v***_***t***,***j***_ is perpendicular to LOS. **(b)** Geometry of finding the subtended angle *α*_*j*_. Edge rays are denoted with green lines. Edge rays pass at two corners of the neighboring agent, and the segment between those points we define as the ‘effective edge’ ***d*** (here, ***d***_***A***_ for neighbor *A*, ***d***_***B***_ for neighbor *B*, etc.). Depending on the relative orientation of the neighbor with respect to the focal agent, the effective edge may be either the neighbor’s diagonal (see neighbor *A*), its shorter side (neighbor *B*), or its longer side (neighbor *C*).

Taking a reductionist approach, we only assume the single omni-directional sensor can measure subtended angles and—over multiple frames taken in successive time—angular displacements of tracked objects (Figs. 2a and 2b). It does not measure the orientation or heading of the observed neighbor, since identifying orientation requires depth perception ability. As a result, inferring inter-agent distances from the angular projection of a neighbor is generally impossible, as different distances can produce equal projections (Fig. 2b). This also raises a challenge for estimating the velocity vector ***v***_***j***_ for neighbor *j*, as different actual velocity vectors can be projected to identical observed angular displacements (Fig. 2c; see also [35]). The elongated morphology of the agents is a crucial factor in the accuracy of this process: when agents are circular, the projected subtended angle allows for accurate estimation of the distance, and thus to the precise knowledge of displacements and velocity (see Figure 2d, and an extended discussion in Section 5).

**Fig 2.**
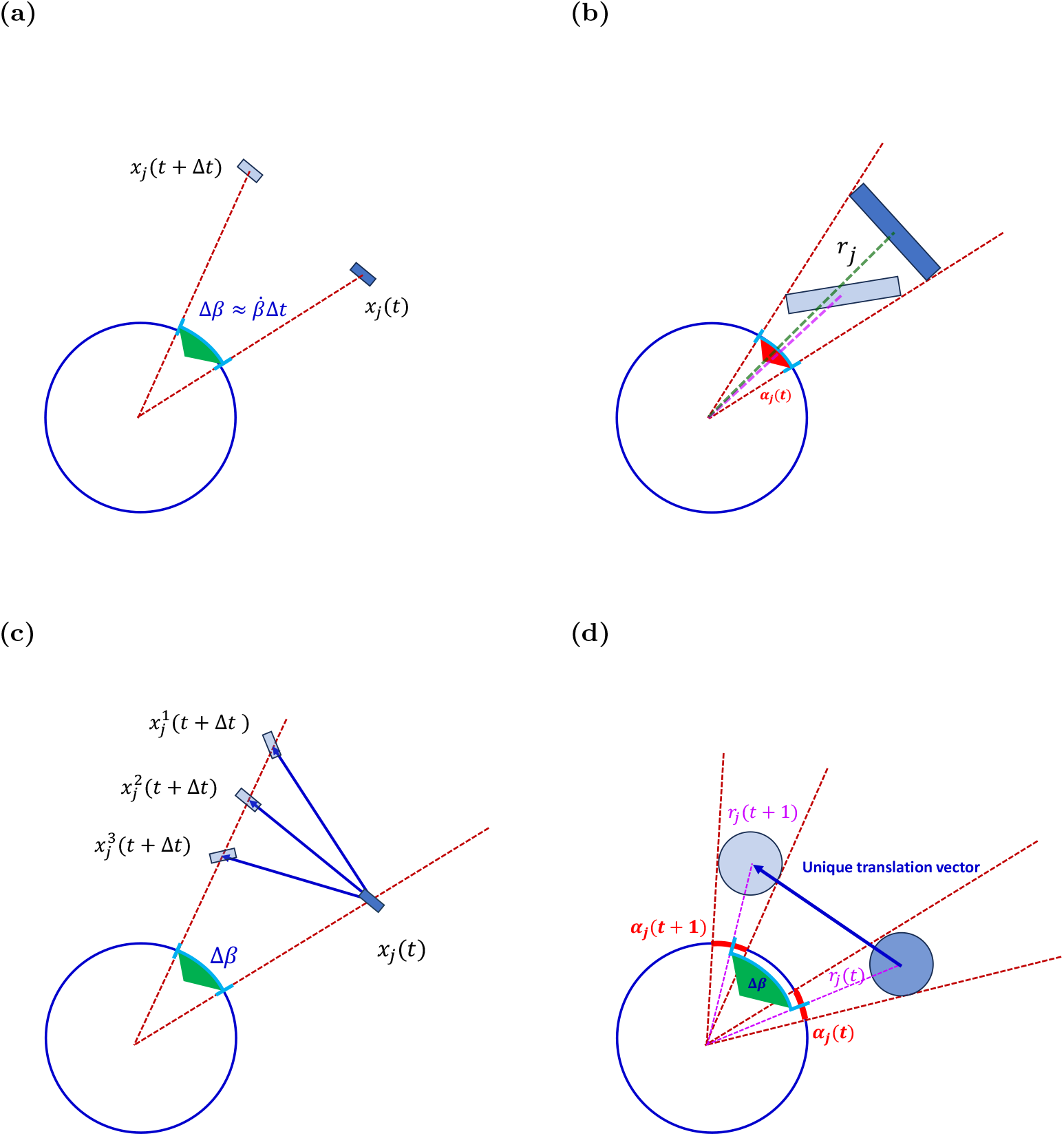
Sensing pointing angles and angular displacements. The blue circle represents an idealized 360-degree visual sensor of the focal agent. Positions at time *t* are marked by *x*(*t*). The elongated shape of the neighbor agent leads to ambiguity in computing its kinematic parameters when using solely angular data. In contrast, for circular agent morphology, the angular data are sufficient to extract complete and exact kinematic data of the neighbor. **(a)** Angular velocity 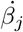 is computed from Δ*β*_*j*_, i.e., from the change of the LOS direction. **(b)** Distance *r*_*j*_ to the neighbor j is estimated from the angle *α*_*j*_ subtended by the neighbor, using Eq. 5. Different distances to the neighbor (green and purple lines) can have the same subtended angle *α*_*j*_ due to the different orientations of the neighbor with respect to the LOS. **(c)** A related source of ambiguity lies in the impossibility of computing the components of the neighbor’s velocity accurately when using only angular information from a single visual sensor. As shown: many different endpoints produce equal Δ*β*. **(d)** In contrast: when agents are circular, the angular information *α*_*j*_ and Δ*β*_*j*_ suffices for an exact computation of distance and velocity, because the distance *r*_*j*_ is uniquely obtained from *α*_*j*_ alone.

#### Estimating neighbors radial and tangential velocities

We start the estimation of neighbor’s *j* velocity ***v***_***j***_ by separately estimating its two components ***v***_***j***,***r***_ (radial velocity) and ***v***_***j***,***t***_ (tangential velocity) (illustrated in Fig. 1a). Both components are estimated on the basis of the instantaneous vectorial distance ***r***_***j***_. We make two assumptions in computing this estimate, with respect to the orientation and size of the observed neighbor, as discussed below.

First, since the orientation of the neighbor is unknown to the observer, we use a simplifying assumption that the neighbor’s effective edge (*d*, in Fig. 1b) is *perpendicular* to the LOS. Commonly, this effective edge would be the diagonal of the observed rectangle (neighbor **A** in Fig. 1b), as observing the rectangle edges occurs only in rare cases of perfectly parallel or head-on motion. The triangle comprised of the focal agent’s COM_*o*_ and the two vertices of the effective edge ***d*** (see Figure 1a) is then taken to be equilateral (see details in A.1). Under this assumption, the LOS constitutes both median and altitude to the effective edge, and a bisector of the subtended angle *α*_*j*_ (see Fig A.1), and therefore, the scalar distance *r*_*j*_ is given by

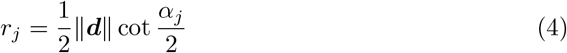

and the vectorial distance ***r***_***j***_ is given by

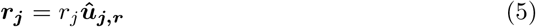

where ***û***_***j***,***r***_ is the unit vector pointing toward the neighbor *j* along the LOS to it (see Figure 1a, and Appendix A.1).

The distance estimation is based on a second assumption, as also made by other researchers [63–65] that animals can possess knowledge of the *typical* size of its conspecifics, especially in homogeneous swarms. In our case, this translates into an assumption that the effective edge *∥****d****∥* used in Eq. 4 is a known *constant* for the agents. Combining this constant *d* with the angle (*α*_*j*_), one can estimate the distance vector (Eq. (5)). This estimate has been used in earlier studies in the context of loom calculations [61, 63, 64, 66].

We emphasized that *r*_*j*_, as given by Eq. 4 is an inaccurate estimate of the actual distance to the neighbor *j*, because it is based on the assumption that the effective edge ***d*** is always perpendicular to the LOS, which is not true in general, and is of given constant length (typical of conspecifics). In reality, the effective edge depends on the specific instantaneous orientation of the observed neighbor, as shown in 1b, and on its actual size.

Relying on the two assumptions above, the radial velocity is computed by differentiating Eq. (5) with respect to time *t*,

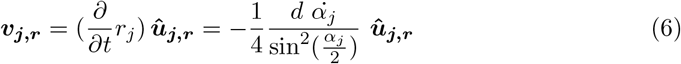

where 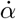 denotes the time derivative of the subtended angle. Expressing 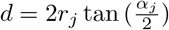 from Eq. (4) and substituting into Eq. (6) results in the radial velocity ***v***_***j***,***r***_, (the derivation is detailed in Appendix A.2):

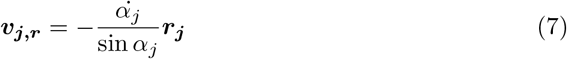

The negative sign means that when the subtended angle increases, the velocity of the neighbor is towards the focal agent, and vice versa; see Figure 1a for the intuition.

While the radial velocity is estimated only from the projected subtended angle and its rate of change, the tangential velocity requires additional components: the bearing angle *β* (which is generally known), and its derivative over time 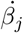 (also known as the instantaneous angular velocity):

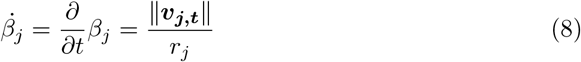

from which we can deduce

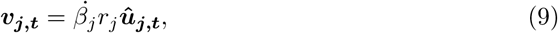

where ***û***_***j***,***t***_ is the unit vector of the tangential direction, i.e., perpendicular to the radial unit vector ***û***_***j***,***r***_.

Combining the two components, we obtain the full velocity vector of neighbor *j*, ***v***_***j***_ = ***v***_***j***,***r***_ + ***v***_***j***,***t***_. This process is repeated for all the neighbors, and the mean ***v***_***j***_, (***v***_***desired***_ of the focal agent) is computed by the formula in Eq. 3.

We emphasize that this is a baseline model. It assumes that all the neighbors are fully visible and does not account for possible obstructions of sight. In other words, the agents are presumed to be *transparent*, in the sense that they do not occlude more distant neighbors. Because this assumption clearly ignores fundamental limitations of visual perception in nature or in robots, we explore general strategies to address it in the next section.

### 2.2 Addressing Occlusions: Three Approaches

Occlusions present an inherent challenge to the use of visual modality in both natural and synthetic agents. Flocking swarms, whether natural or artificial, are often dense [67]. Conspecifics located closer to the observing animal are inevitably blocking, partially or entirely, the animals standing behind them (Fig. 3).

**Fig 3.**
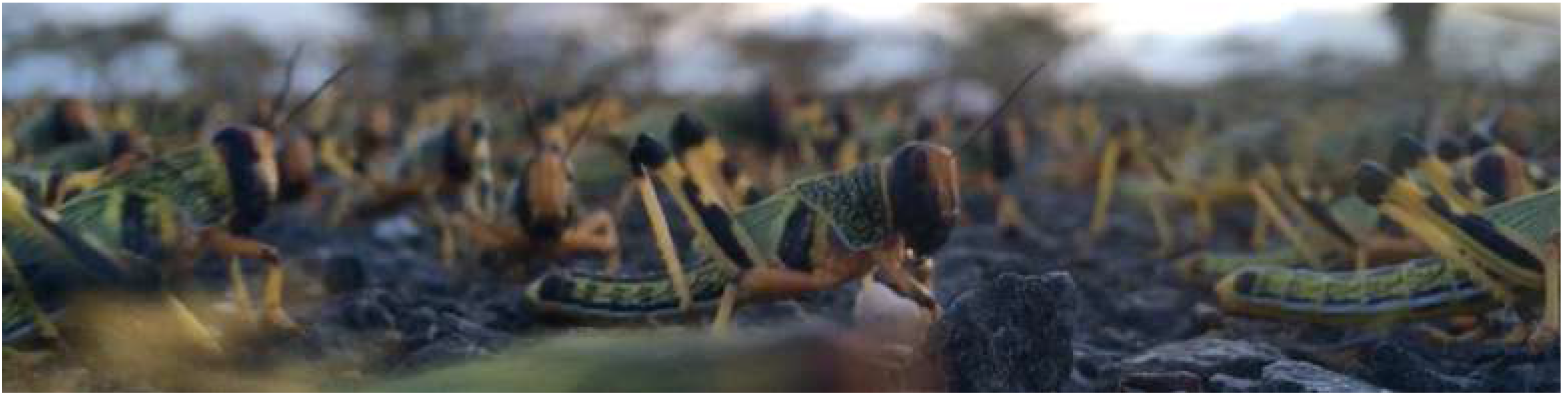
The visual social environment from the perspective of the individual locust in a swarm. Picture credit: Inga Petelski, Kenya, 2020.

Complete and partial occlusion of neighbors not only reduces the information available to the focal agent but can also introduce large estimation errors. Neighbors that are completely occluded are not taken into account in the collective motion model. *Partially* -occluded neighbors introduce errors, as the projected area of their subtended angle, used as a proxy for distance, is smaller than it should be. For example, suppose a neighbor is partially occluded, such that only a small portion of it is observed, and thus it is initially perceived to be distant: if the occluding animal moves to uncover it, its full length will now be revealed, and within a very short time it will be seen as being close, implying high radial velocity towards the observer and a potential collision. The accumulation of such frequent errors may disturb the stability of the swarm.

We posit there are three general strategies that may be applied (illustrated in Fig 4). Suppose the focal agent may be able to recognize peers and thus differentiate between entirely-visible individuals and parts (partially-occluded individuals that are not recognized as conspecifics). This allows it to ignore partially-visible neighbors (Fig 4a). It may also be able to cognitively extrapolate parts to a whole, inferring the position and orientation of the partially-occluded peer from its visible parts (Fig 4b). Alternatively, without being able to recognize peers, the focal agent may still be able to perceive any visible part of a neighbor as a distinct whole individual. These different strategies place very different requirements on the cognitive-computational processes of visual perception in the focal agent, as discussed in detail below.

**Fig 4.**
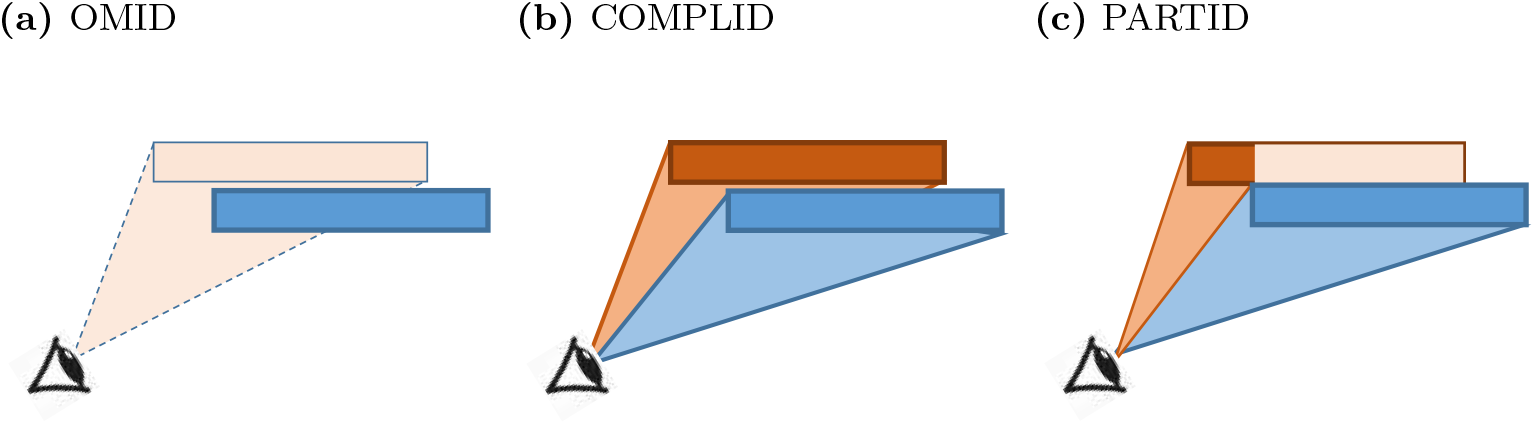
Occlusion methods. **(a)** OMID – partially occluded neighbor (orange) is omitted from the field of view. **(b)** COMPLID – orange neighbor is completed from the seen segment. **(c)** PARTID – partially seen segment is regarded as a neighbor.

#### Approach 1: Omission of the Occluded (OMID)

The first approach disregards any visual information originating in partially occluded agents (see Fig 4a). This requires the animal to possess a dedicated *peer recognition* mechanism, i.e., to be able to recognize fully-imaged conspecifics (and ignore anything else). Mechanisms of selective attention in visual perception are known to exist in humans and are achieved in the human brain in multiple stages of perception [68, 69]. Neurobiological studies have shown the existence of selective attention mechanisms also in insects’ visual processes [70, 71].

However, it is not known whether locust visual perception mechanisms are able to recognize peers. Experiments reported by Bleichman et al. [36] have shown that an individual locust responds by walking when exposed to visual images composed of randomly-moving dots that are projected via computer screens to both eyes. As the dots are positioned randomly and do not mimic the shape or the colors of locust nymphs, these results seem to indicate that the motion is triggered in the individual devoid of any dedicated peer recognition mechanism. Nevertheless, such visual processing may, in principle, be applied during collective motion and constitute a plausible approach that exists in nature.

#### Approach 2: Completion of the Occluded (COMPLID)

In the second approach, partially occluded agents are “completed” as if they are fully visible to the observer. In other words, a neighbor that presents even the smallest visible segment from the focal agent’s perspective would be treated as if no occlusion is present when processing its visually extractable information. COMPLID utilizes *peer recognition* as in OMID, while also requiring that the agents will be able to assess the obscured part of a neighbor (if needed) based on its visible part. This completion assumes an agent’s visual extrapolation that reconstructs neighbors’ outlines using their visible features.

Completing partially visible targets obscured by other objects is a long-studied process in visual perception. The filling-in of details and image regions partially obscured by interceding objects [72, 73] is an established neurophysiological process that gives the organism an ability to identify a complete form based upon observed parts of the contour and is described by the term “visual completion” [74]. This mechanism produces an internal representation called “illusory contour”, which extrapolates the physical stimulus to the full geometrical shape of the object [29, 75, 76]. Visual completion of occluded objects has been shown in varied and phylogenetically distant species: birds, fishes, cephalopods, bees, etc., and is accepted as one of the fundamental components of vision in nature [75, 77, 78].

#### Approach 3: Every Part is a Full Agent (PARTID)

The third approach treats all visual stimuli related to a neighbor as if they represent a full-body conspecific. Contrary to OMID and COMPLID, this approach makes no assumption of *peer recognition* capabilities. Rather, the visual field is divided into segments, with each segment containing the same optical flow vectors. The agent assumes that each segment represents a different neighbor. In other words, any visual information is taken completely at face value without any additional interpretation. Hence, other than the ability to accurately extract optical flow vectors, no further advanced visual perception mechanisms are required. Since the optical flow is essentially the vectorial difference between two consecutive frames and does not consist in any form of object recognition by itself, PARTID would be the least computationally demanding approach if implemented in real life.

However, in this approach, the potential error in the assessment of the environment is the largest, in comparison to OMID and COMPLID, since partially occluded agents occupy less area on the visual field, which translates to a significantly larger distance estimation. The same applies to velocity estimations, which are tightly dependent on the distance. Although in this approach, an agent does not possess with object recognition abilities, it is assumed that the observable parameters 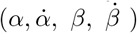 are still fully available and extractable. As noted, this approach requires relatively limited visual processing and is easier to implement in robotic systems.

PARTID takes its inspiration from biological mechanisms, in which an organism performs an action based on visual stimuli originating from an object that is not recognized. For example, locusts possess a pair of visually-sensitive neurons that encode looming stimuli and cause the locust to produce escape behaviors [61]. The visual stimuli affect the behavior of the individual directly and without passing through object recognition mechanisms [36].

### 2.3 Summary

We summarize the different mechanisms introduced in this section. First, we derived estimates for the velocities of visible neighbors, such that these velocity vectors can be aggregated in a Viscek flocking mechanisms for determining individual velocity at any given moment. These estimates rely on assumptions with respect to the background knowledge available to the individual (the typical size of conspecifics), as well as on the orientation of the observed agents (parallel to the line of sight). We refer to this base reductionist model as the *principal* model.

We then discuss strategies for addressing occlusions, which can further degrade the accuracy of estimated velocities. All three strategies ignore completely occluded agents (unlike the principal model) but differ in how they treat partially occluded neighbors. They are summarized in Table 1 below.

**Table 1.** A summary of the differences and similarities between the different reductionist models. Rows present different occlusion conditions with respect to the neighbor in question. Columns contrast the various models in how they respond to these conditions. See also Figure 4.

| Neighbor visible? | Principal | OMID | COMPLID | PARTID |
| --- | --- | --- | --- | --- |
| Completely occluded | Fully visible | Ignored | Ignored | Ignored |
| Not occluded | Fully visible | Fully visible | Fully visible | Fully visible |
| Partially-occluded | Fully visible | Ignored | Fully visible | Part is neighbor |

## 3 Methods

In order to evaluate swarm behavior using different occlusion-handling approaches, we developed a two-dimensional (2D) collective motion simulator based on a basic simulation engine [79] (see Fig 5). The agents’ movement in two dimensions is simulated by updating their coordinates at each iteration in accordance with their current velocities and the position update control laws presented in Section 2.1. The location and orientation of each rectangular agent are computed from the coordinates of its COM. It is assumed in our model that velocity heading is always along the long axis of the body. The velocity magnitude can vary between 0 and a fixed maximal speed value *v*_*max*_, i.e., the agents can accelerate up to a maximal speed. Together with the steering parameter *η*, this reduces the sharpness of turns and accelerations in the agents’ motions.

**Fig 5.**
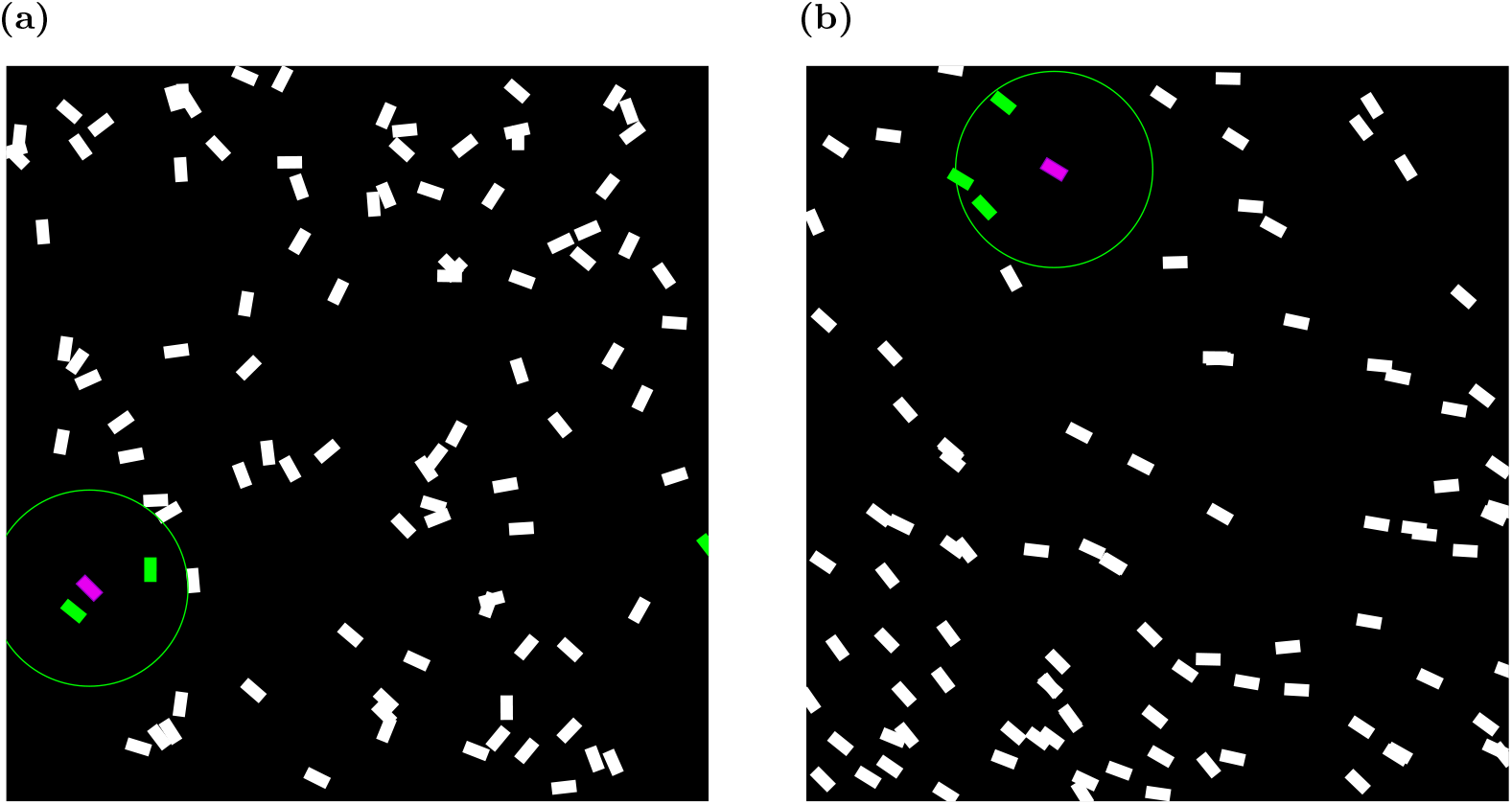
Simulator snapshots. **(a)** Toroidal arena snapshot at *t* = 10[*frames*]. Agents are initialized at random positions and random velocities. The purple-colored agent is an arbitrarily marked focal agent with its respective neighbors colored green. **(b)** Toroidal arena snapshot at *t* = 2000[*frames*]. An apparent flocking behavior is displayed, with all agents moving roughly in a single direction.

**Fig 6.**
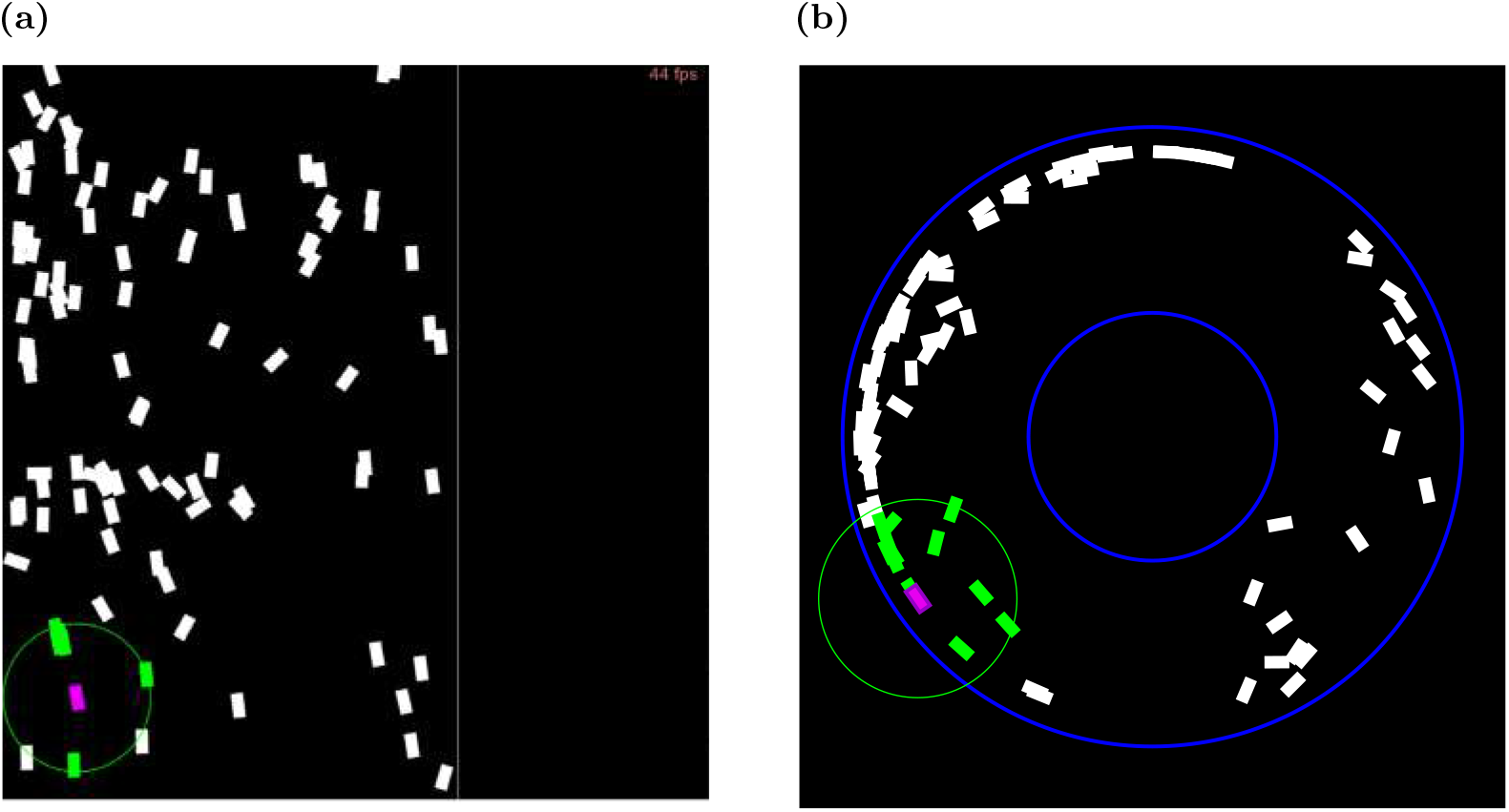
**(a)** Snapshot of corridor arena. The vertical boundaries are repelling, while the horizontal ones are periodic. **(b)** Ring arena snapshot.

The agent’s motion decisions are based on the neighbors’ velocities. These velocities, in turn, are derived from the angular measurements of each perceived neighbor: their subtended angle and the angular velocity. These inputs serve as the agent’s subjective perception.

### 3.1 Simulating Perception

We compare the emergent collective motion resulting from the different occlusion-handling approaches. The perception of each agent is simulated. The exact values stored in the simulation, are used as the basis for emulated perceptual processes, and the effects of occlusions. Each simulated focal agent is given the information it would have perceived in the 2D environment, in the principal model, and in the three occlusion-handling strategies.

#### Simulating the Principal Model

The *α* angle is calculated using the neighbor’s vertices of the edge that subtends the largest angle on the agent, regardless of occlusion. The angle between the two vectors pointing from the focal agent’s COM to the respective vertices equals *α*. The *β* angle is simply the angle between the focal agent’s velocity vector and the neighbor’s COM, again regardless of occlusions. The focal agent receives visual parameters of all the neighbors, *including those that are completely occluded by others*.

#### Simulating OMID

All completely occluded or partially occluded neighbors are ignored. The effective *α* and *β* for each completely-visible neighbor are taken from the subtended angle as before, and only those are used in computing *v*_*desired*_.

#### Simulating COMPLID

We simulate this capacity by means of calculation, taking the same measurements as in the principal model. We then remove from consideration all neighbors fully occluded by others.

#### Simulating PARTID

We iterate over the neighbors, from the closest to the furthest. Each neighbor’s effective edge is calculated and then checked against an array of edges. If a partial overlap occurs with the current edge and one or two of the already checked edges, the effective *α* is calculated using only the non-overlapping segment: that is, the subtended angle from any visible part of a neighbor is taken to be a neighbor, and its center of mass is taken to be the angular midpoint.

### 3.2 Controlled (independent) simulation variables

The simulator enabled control of the many variables. The population size *N* controls the number of agents in the simulated swarm. The body length-to-width ratio determines the elongation of the agent, and thus is assumed-constant effective edge size *d*. The effective range of the sensor, *R* is measured in body lengths ([BL] units), and determines the range within which the agent is able to perceive neighbors, without occlusions. The steering parameter *η* sets the weight of *v*_*desired*_ relative to the current velocity (*v*_*i*_) of an agent. *v*_*max*_, which caps the maximal speed attainable by agents and was arbitrarily set to 1[BL]/frame.

We utilized different areas (*arenas*) in the simulation experiments: a square arena with periodic boundaries, an infinite corridor where only one axis has periodic boundaries and a circular arena (with no periodic boundaries). Where a period boundary occurs, once an agent’s COM passes the maximal/minimal coordinates or the X/Y axes, it reappears on the other side respectively. Where a non-periodic bound is reached by an agent, it is repelled with varying repelling force, depending on the size of the radial velocity component (relative to the arena center), i.e., an agent traveling to the external circular boundary will be repelled from with force proportional to the size of the agent’s radial velocity component. This was designed to mimic the behavior of live locusts that align themselves to the ring walls in order to avoid possible collision with them [80].

### 3.3 Measured (dependent) simulation outcome: Flocking Order

The ideal flocking is a situation in which all agents are synchronously moving in the same direction. Over the years, various measures of order have been proposed and utilized in different settings. As we essentially extend the Vicsek-based collective motion model to account for visual perception, we chose the *polarization* measure of order, denoted *ϕ* and used in other investigations of Vicsek-based collective motion [14, 20, 81–83]. It is defined by

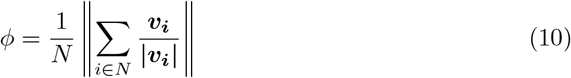

where *N* is the population size, and ***v***_***i***_, ***∥v***_***i***_***∥*** correspond to the velocity and speed (resp.) of agent *i. ϕ* measures the degree of global alignment by averaging the normalized velocities of the agents (i.e., headings). It is a scalar value representing at any given time the degree of global order in the system. For a random disordered group state, *ϕ* is approximately 0, while for a fully ordered flock, with all agents moving with an identical heading, it approaches a value of 1.

## 4 Results

Two sets of experiments were conducted to evaluate the presented approaches. The first set, presented in Section 4.1, uses the principal model to set baseline parameter ranges for various controlled settings. The second set of experiments, presented in Section 4.2, then uses the established parameters to contrast the performance of the three occlusion strategies alongside the principal model in different arenas, whose geometry and bound periodicity are varied.

Unless otherwise stated, the experiments comprised 50 independent trials, each with its own randomized initial conditions (individual velocities, including headings), such that the swarm was unordered (*ϕ* close to 0). The figures present the mean over the 50 trials, with error bars (or shaded envelope around the solid lines) showing margins defined by the standard error. This enables the distinction of significant differences between different models.

We remind the reader that we follow others in using the polarization order *ϕ* as the primary measure of order in the flocking system, in which the range is no-order to high order, 0≤ *ϕ ≤*1. We typically present two types of graphs. The first shows the *evolution of the order parameter ϕ over time*, measured for simulation time (frames) *t* = 1 … 3000, in which we terminate the simulation for practical reasons. It is frequently but not always, clear that *ϕ* converges to a value by the end of the simulation, which can indicate that the swarm is converging to an ordered state. However, this is not always the case. Sometimes *ϕ* increases monotonically (which may indicate convergence at a slower rate), and other times, it remains close to 0. It is, therefore, useful to also display a second type of graph, which shows the *long-term* (end-of-simulation) *ϕ*, obtained at a fixed time of *t* = 3000 [simulation frames]. This allows quick determination of the convergence of the swarm within the allotted time frame but does not indicate a lack of convergence should the simulation be allowed to run indefinitely.

### 4.1 Flocking using the Principal Model: Baselines

We begin by testing the principal model in a toroidal arena, varying independent simulation variables chosen in accordance with observed locust characteristics. The goal is to establish baseline responses to various settings, such as the visual range *R*, the steering parameter *η*, etc. As simulation measurements are artificial, we use a standard length unit, [BL], which is the agent’s default body length, with a length-to-width ratio of 3. For the experiments reported in this section, we used an arena of size 20 *×* 20 [*BL*^2^].

#### 4.1.1 Determination of Steering Parameter

The first experiment sought to determine an appropriate steering parameter, *η* empirically. Initial settings were based on observations of locust marching bands: a population size *N* = 120 within the arena resembles reported marching locust density in nature [67] (see below for other values of *N*). Similarly, the sensing radius *R* = 3 body lengths ([BL]) was set according to empirical observations of locust nymphs not reacting to visual stimuli located farther than 2–3 [BL] [26]. The agent elongation (body length ratio) was set to 3 (i.e., agent length is three times its width; see Section 4.1.3 for discussion).

We experimented with different values of the steering parameter *η*. Fig. 7a shows the mean order measure *ϕ* as it changes over time, measured in simulation frames (*t* = 1 … 3000), for four values of *η*. It can be seen that smaller values of *η* cause the swarm to converge towards a higher order, while larger values do not.

**Fig 7.**
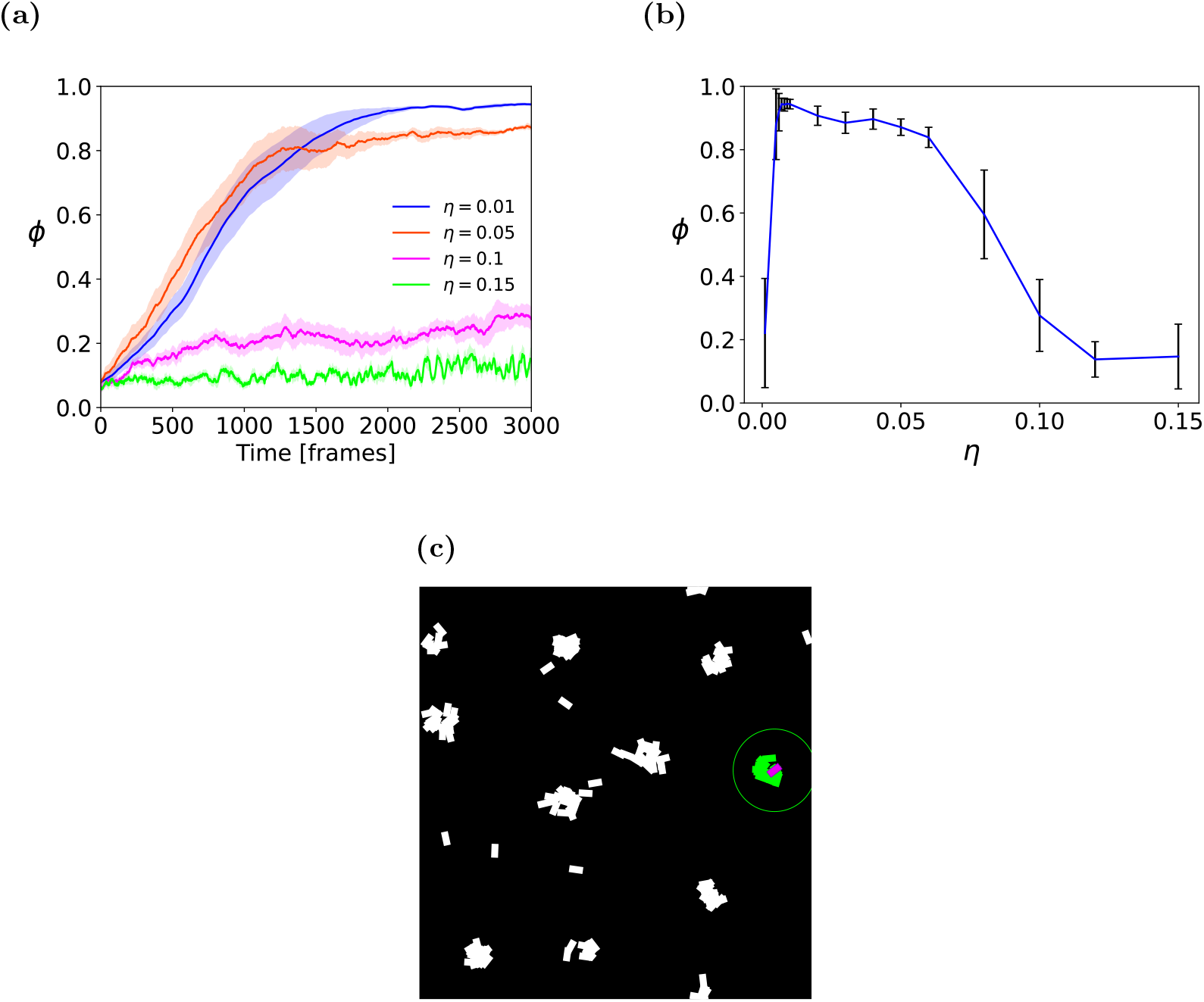
Steering parameter sensitivity analysis, over 10 independent runs. **(a)** Mean order (*ϕ*) for different *η, t* = 1 3000.The solid line shows the mean order parameter of the swarm for each *t*, with standard error margins shown in the envelope. **(b)** Long-term (*t* = 3000) mean order (*ϕ*) for varying *η* values. **(c)** *Cluster* pattern of agents moving under high *η* values.

Fig. 7b examines a more extensive set of *η* values in terms of the order measurement at time *t* = 3000. It can be seen that a value of *η* = 0.01 yields the maximal value of the order parameter, approaching 1. This is where the swarm is nearly fully aligned. Notably, no convergence occurs for smaller values, meaning that the agents are apathetic to the environment and retain their original heading directions. In contrast, a significant drop can be seen in the order parameter magnitude for large *η* values, i.e., the agents’ convergence fails due to over-sensitivity to the external steering parameter. Further analysis of these large *η* values is provided in Fig 7c. It shows that agents aggregate in small and tight clusters and constantly change their headings, unable to reach either a local or a global uniform moving direction. Based on these findings, we fixed the steering-parameter factor parameter as *η* = 0.01 for the rest of the experiments reported in this study.

#### 4.1.2 Influence of Vision Radius *R*

A second series of experiments examined the role of the visual sensory range (distance-wise). Initial settings, based on empirical observations of locusts, have set the range *R* at 3[BL]. In this subsection, we examine other values.

Figures 8a–8b, present the evolution of order *ϕ* over time, and its long-term values, for different visual ranges 0.67 ≤*R ≤*3.67 [BL] for a swarm of size N=100. Figure 8a shows the order developing over time for different values. Figure 8b shows long-term mean values of *ϕ* at the end of the simulation (*t* = 3000). As expected, for *R* smaller than 1[BL], the progress toward the ordered group state is weak and very slow. The reason for that is: for such short-range of visibility, most neighbors are unobserved. Thus, vision does not provide sufficient information about the neighbors to the focal agent. For larger values of *R*, the long-term *ϕ* slightly increases with larger radii. Interestingly, lab experiments and observations of locusts have estimated their visual range to be 2–3[BL]. For the remainder of the experiments, we set *R* = 2.67[*BL*].

**Fig 8.**
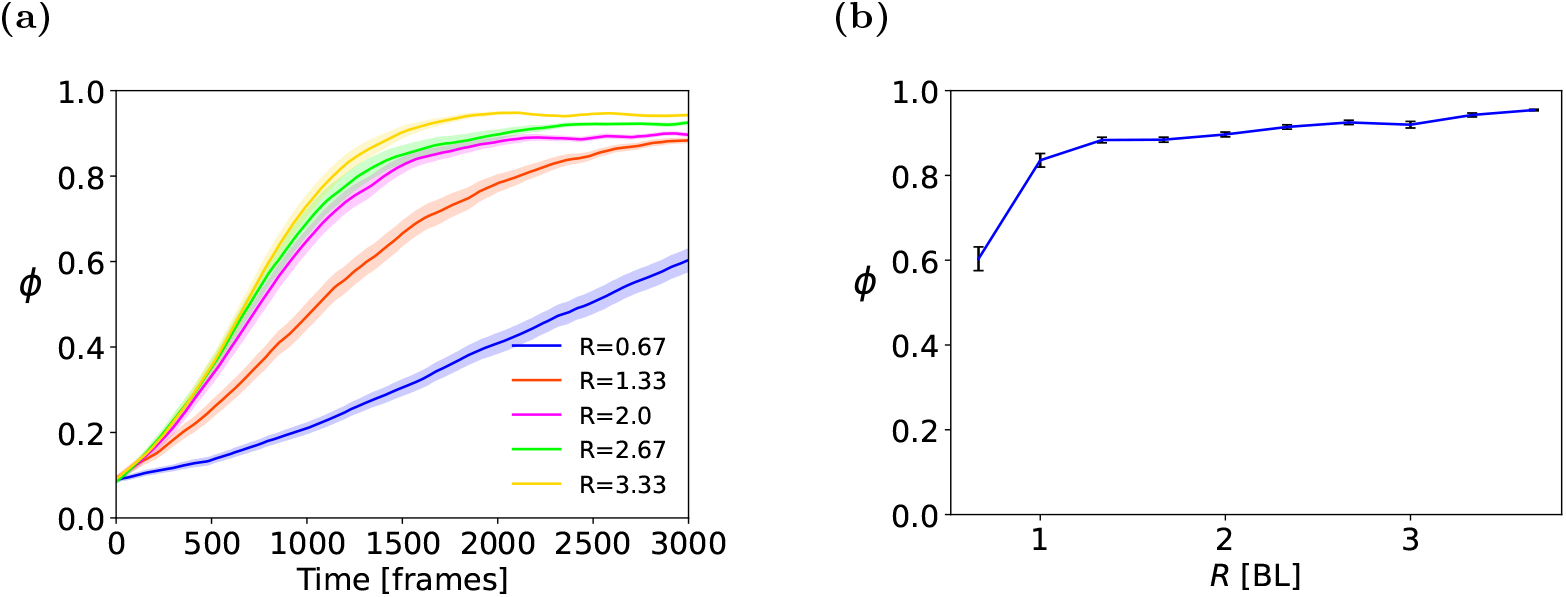
**(a)** Time-dependent and **(b)** long term sensitivity analysis for visual range *R*, measured in body lengths [BL], in the Torus arena. Means and standard errors shown for 50 trials.

#### 4.1.3 Other influences on flocking order

Interconnections clearly exist between the different parameters. For instance, it is possible that for visual ranges that dramatically differ from those we used in our experiments, different values of *η* will yield different results. Similarly, varying the type of environment used can influence the rate of convergence or even its existence. As we sought to explore the model as inspired by nature (in particular, locust), we set out to experiment with settings in ranges that approximate locust swarms and used the toroidal arena and principal model, as we believe these to be the least constraining, and least sensitive to parameters that are external to the model itself.

The population size *N* is a clear factor regarding the emergence of order, as varying *N* while maintaining a fixed arena area (or inversely, varying the arena size while maintaining a fixed value of *N*) impacts the swarm density. This in turn influences the likelihood of occlusions, the ability—given limits on *R*—to observe neighbors, etc. In the different arenas we set values of *N* that we had experimentally determined to be informative, in that they reveal differences between the different strategies. Appendix A.3 shows how this procedure was carried out for the torus arena. We took similar steps to determine *N* in the other arenas.

We now turn to discussing the body length ratio, which measures the elongation of the agent. We used a length-to-width ratio of 3 [BL] unless otherwise noted, as this approximates the observed dimensions of typical locust nymphs in our laboratory, which inspired this research. This is a critical issue, as some existing models of vision-based flocking use non-elongated (circular) agents. While locusts, and many other swarming species, are clearly elongated, it is important to establish whether the elongation (as measured by the body length ratio) influences the results. Otherwise, non-elongated agents—circular or squares—could equally serve as a model for locusts or other elongated agents.

Appendix A.4 provides an empirical exploration of the influence of the length-to-width ratio on convergence, in various environments, and in all flocking models (principal, OMID, COMPLID, PARTID). Briefly, the results show that convergence to an ordered state is highly sensitive to the length-to-width ratio, and thus setting its value to model locust body dimensions is critical. As these results complement the main results for the occlusion-handling strategies that we report below, we advise the reader to examine them after the main body of results is presented. We also address this issue in the Discussion (Section 5).

### 4.2 Comparison of the three occlusion strategies and the principal model

Having established the baseline parameter and experiment settings, we now turn to investigate the emerging order *ϕ* of swarms, utilizing different strategies. The three occlusion-handling strategies are evaluated in comparison with the *principal* model (which does not account for occlusions). A summary of the commonalities and differences between the models is provided in Table 1.

#### 4.2.1 Experiments in the Torus Arena

We begin with the experiments in the *Torus* arena, which we had utilized (above) for establishing the baseline parameter values. Figure 9 shows the evolution of the order *ϕ* over time for all four strategies. The graphs show the mean order parameter for each point in time. Three population sizes of *N* = 60, 120, 180 are shown; in all experiments *R* = 3[BL], *η* = 0.01, and length-to-width ratio is 3.

**Fig 9.**
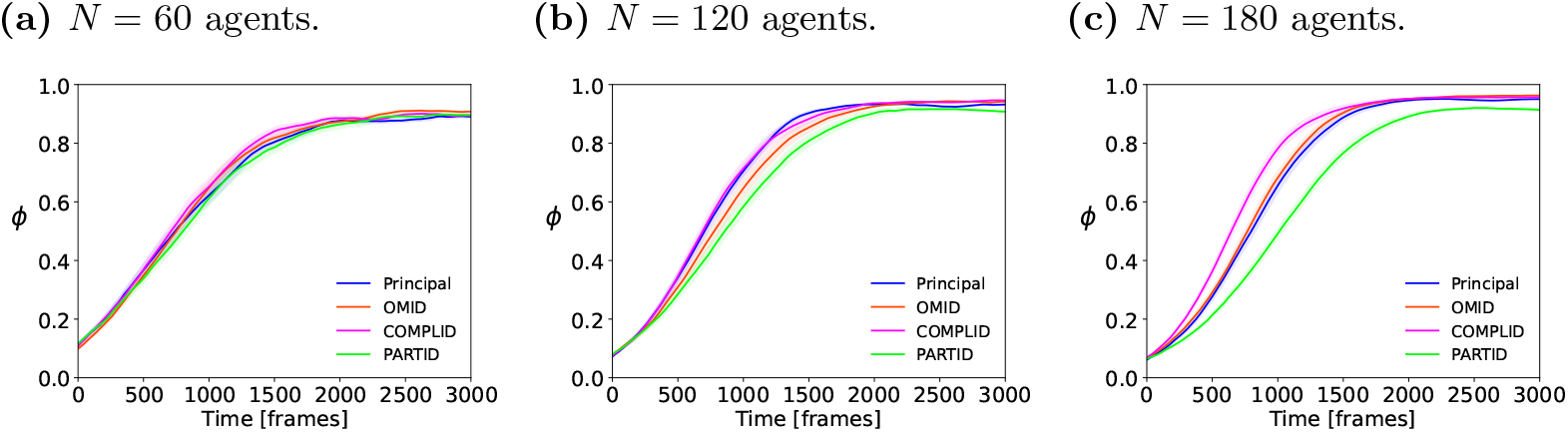
*ϕ* evolving over time *t* = 1 … 3000, for different strategies, in the torus arena. Plots show the mean order parameter of the swarm at each simulation frame, with standard error margin for different population sizes, over 50 independent trials. Larger *N* generally leads to a slightly steeper transition to a flocked state.

At higher densities (shown in Fig 9c) the rate of convergence of all three perceptive approaches lags behind the principal model. At higher densities, rates of convergence become steeper. At the same time, the long-term order parameter remains very close for all the methods, and even different densities. Finally, the ordering of the rates of convergence at 180 indicates that COMPLID converges faster than OMID. Completing parts of neighbors, rather than omitting them, leads to an effectively larger number of neighbors, which leads to stronger alignment.

Figure 10 complements Figure 9 above. It shows the long-term mean order at the end of the simulation *t* = 3000. It is evident that all three occlusion approaches, alongside the original model, reach similar long-term order-parameter values (*ϕ ∼*0.9), indicating their reaching i.e., similar degrees of ordered flocking. That said, when we consider the result of PARTID at *N* = 180, and also examine its behavior in Figures 9b and 9c, we see that PARTID has a slower rate of convergence, and slightly lower long-term order (note the separation defined by the standard error bars for PARTID when *N* = 180, in Figure 10). This can be interpreted as additional evidence that PARTID is may generate excessively noisy perception.

**Fig 10.**
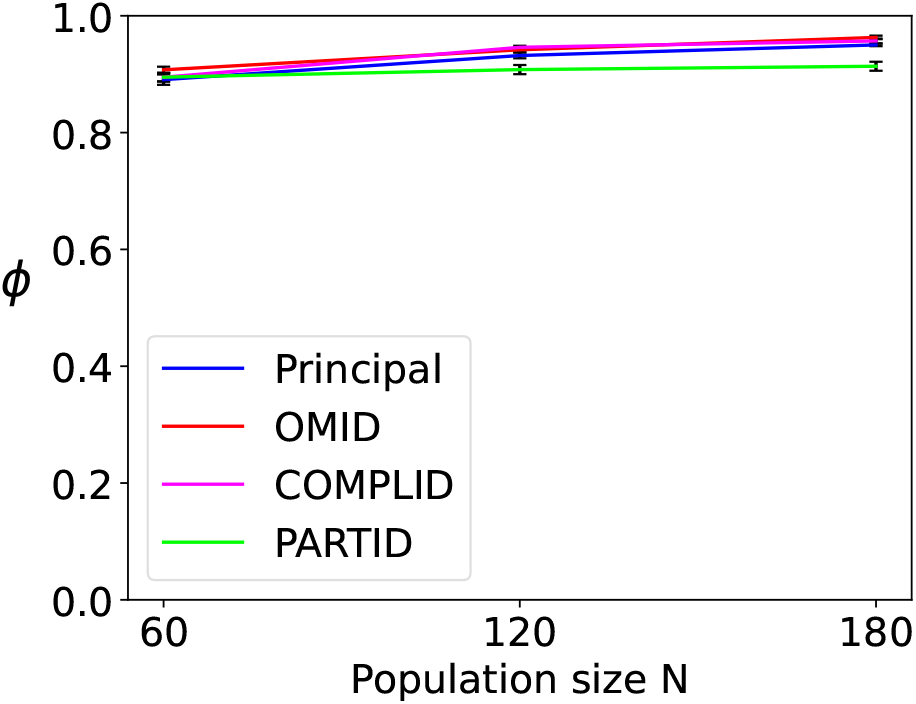
Long-term order of the four strategies: principal, OMID, COMPLID, PARTID. The plots shows the mean (and standard error) long-term order *ϕ* for *t* = 3000, for different *N* values (50 independent trials). Long-term *ϕ* values are practically indistinguishable.

#### 4.2.2 Experiments in Bounded Arenas

The torus arena is fully periodic: agents moving towards an edge are not repealed by it nor blocked. Rather, they move through it to appear on the opposite side of the arena. Likewise, agents close to one edge can visually sense neighbors that are on the “other side of the edge”, i.e., on the opposite side of the arena. While this is a common arena model in theoretical studies of swarms, its abstract nature distances it from the geometrical constraints of realistic environments, which have bounds and obstacles that impose limits on the movement of the agents.

We therefore switched to experiments in the *infinite corridor* (periodic on one side, but not the other) and the *ring* (non-periodic) arenas, described in Section 3. Three versions of each arena type were tested: *wide, intermediate, and narrow*. The geometry of the arenas is characterized by the arena width to single agent body-length, i.e., arena width in in [BL] units. For the infinite corridor, the distance between the periodical boundaries (length) was 20 [BL] for all the experiments. The widths were: 10, 20, 30 [BL] respectively. For the ring arena the radius of the inner circle was 1.66 [BL] and the outer circle radii tested were: 5, 8.33, 11.66 [BL]. In the experiments below, *N* = 100 (experimentally selected, see Appendix A.3).

Figures 11 and 12 present the results for these settings. In Figure 11, subfigures are arranged in columns distinguishing the width (left to right: wide, intermediate, narrow), and the rows distinguish the arena type (top row: corridor, bottom row: ring). Each subfigure presents the mean and standard error from 50 independent simulation runs.

**Fig 11.**
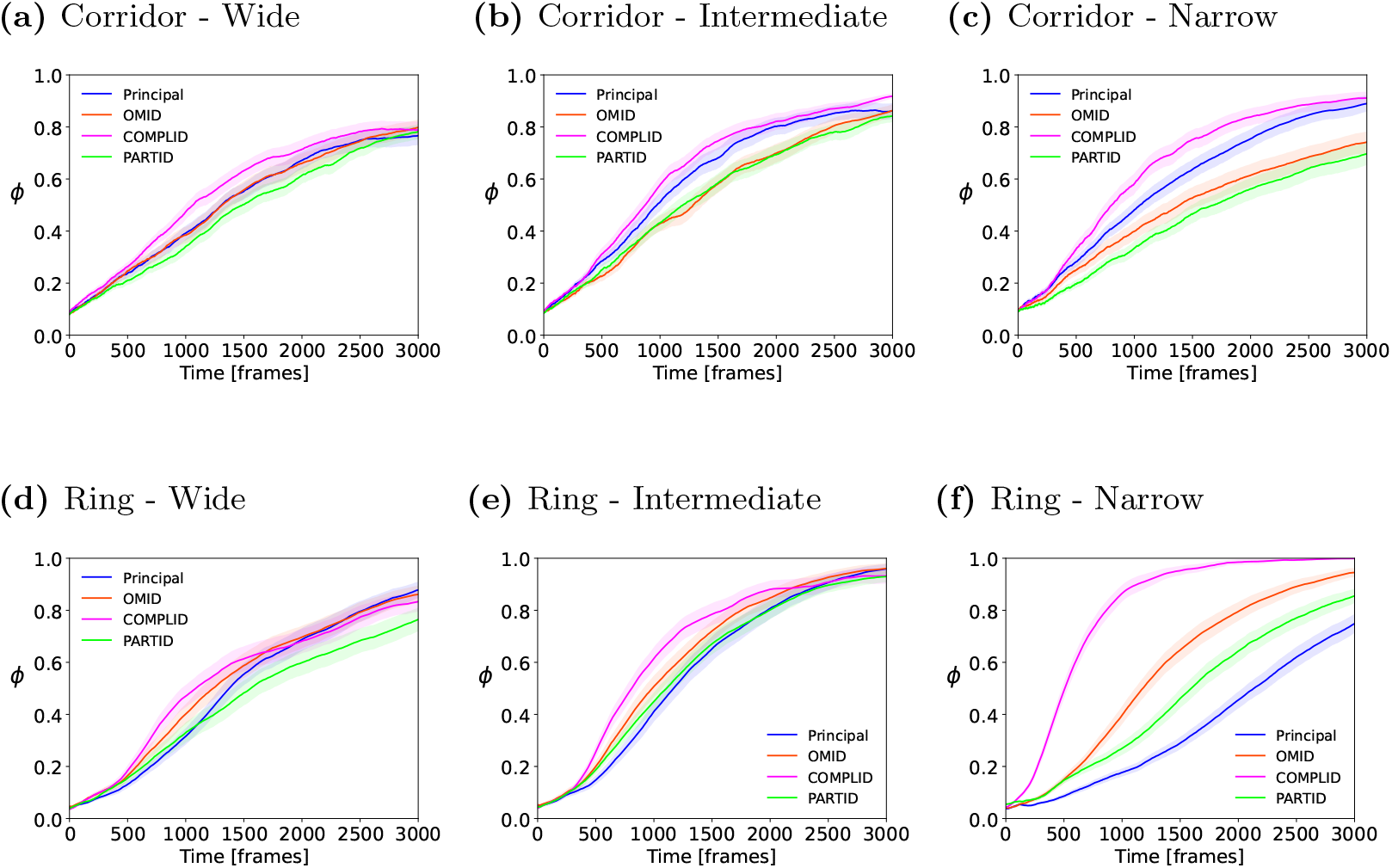
Mean and standard error of the order measure *ϕ*, as it changes over time *t* = 1 … 3000, in 50 trials. **(a)**,**(b)**,**(c)** Infinite corridor arena. Wide / Intermediate / Narrow arena dimensions are 20× 30 / 20 / 10 [BL]). **(d)**,**(e)**,**(f)** Ring arena. Wide / Intermediate / Narrow ring external border radii are 11.66 / 8.33 / 5, respectively. The internal ring border is constant for all three types and equals 2.5.

**Fig 12.**
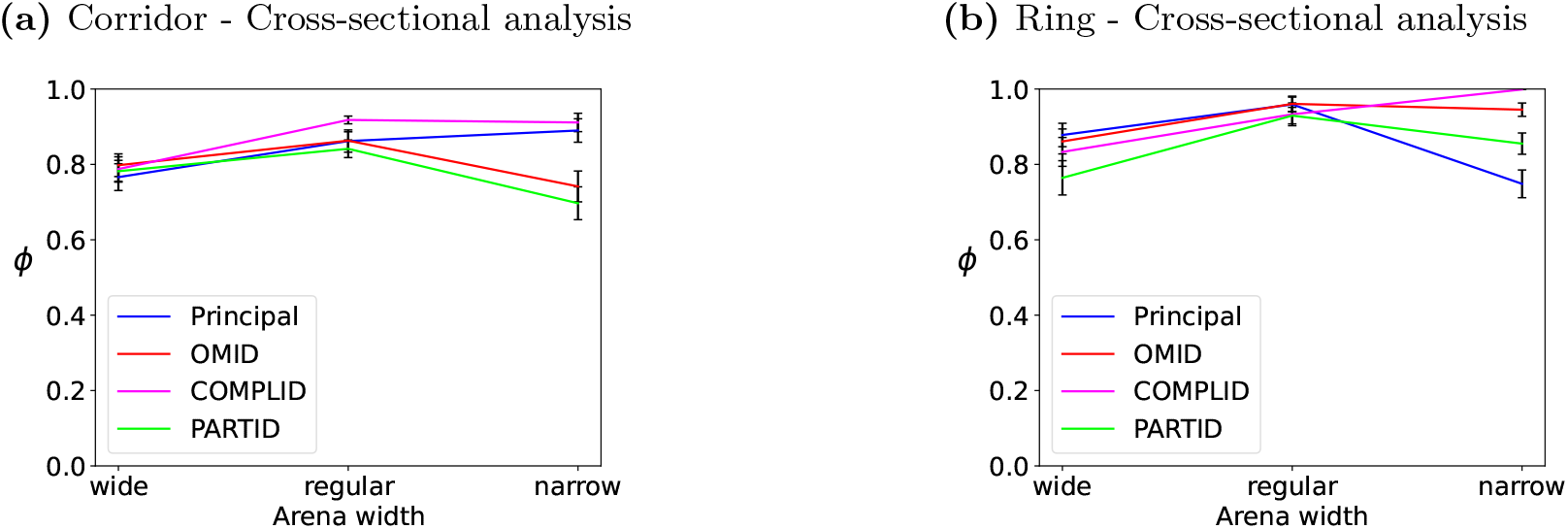
Mean and standard error of the long-term order measure *ϕ*, at *t* = 3000, in 50 trials. In both **(a)**,**(b)**, the horizontal axis marks the width of the arena (wide/intermediate/narrow, as above). **(a)** shows the results for the infinite corridor. **(b)** shows the results for the ring arena.

Each subfigure shows the evolution of the order measure *ϕ* over simulation time *t* = 1 … 3000, for all four strategies. Figure 12 completes the picture, reporting on the long term (*t* = 3000) values of *ϕ* for different strategies, as width decreases (left to right), and density increases.

Figures 11 and 12 combine to show that *narrow* bounded environments (i.e., higher densities, and perturbations caused by bounds pushing the agents back into the arena) cause distinguishable differences in the convergence rate (and success) of the different flocking when utilizing different strategies for handling occlusions. In particular, while all strategies show a rise in the ordering parameter, each strategy’s behavior is distinct. A potential reason for this is the fact that in narrow arenas, interactions with the boundaries are much more frequent. In the corridor, the principal model and the COMPLID strategy converge significantly faster and to a higher long-term value than OMID and PARTID. In the ring, all four strategies are clearly distinguished.

A commonality to both arenas, in all settings, is that the PARTID strategy is generally slower than the others (with the possible exception of wide corridor and intermediate ring). We believe that given that PARTID is a-priori more likely to suffer from noise in the observations, the geometrical bounds, which repel or push agents into the arena (sudden orientation changes) are particularly detrimental to convergence when PARTID is used as a strategy.

## 5 Discussion

The reductionist approach we have taken in this study is intended to shed light on the *necessary* (minimal) mechanisms that can generate an ordered collective motion based on visual processing. This goal stands in contrast to the clear variety of *sufficient* mechanisms that can be used at a computational (cognitive complexity) cost and/or mechanical-physiological requirements. Even when disregarding energy- and computation-hungry sensors and processes used in robots (e.g., LIDAR sensors and associated processing [84]), distance estimates could still be generated from visual information in a number of ways, albeit demanding more capabilities from the agent, compared to the approach we have taken here. For example, stereoscopic vision is a well-understood mechanism for reliable distance estimation [32] in both natural and artificial systems. However, the requirement for overlapping fields of view of each eye (camera) narrows the perceived angle. While multi-lateral distance estimation is required for most traditional models of flocking, quite literally, carrying out the stereoscopic vision in a backward direction would require an additional pair of eyes at the back of the agent’s head [55].

We introduced a non-stereoscopic (monocular) vision-based flocking model for elongated agents in two-dimensions. We considered agents whose motion and perception are situated in two-dimensional flat worlds. The goal was to explore the generation of ordered collective motion with the bare minimum of information. This model employs geometrical aspects of vision, such as subtended visual angle, observable angular velocity, and other derived parameters, but does not otherwise rely on complex visual processes. Unlike many existing models, the model we introduce does not assume direct measurement of inter-agent distances or velocities. Rather, we infer distances and velocities from observed angles and their rates of change. Such measurements are biologically-plausible in non-stereoscopic vision, both in nature and in the robotics lab. Below, we highlight specific issues and questions raised by the results presented above vis-à-vis biological plausibility and other models of vision-based collective movement.

## 5 The Plausibility of Different Strategies for Handling Occlusions

We tested and compared three different general strategies for addressing occlusions in different arenas:

- The first strategy (COMPLID) completes the outline of a partially hidden neighbor. Such abilities are present in various species, and there is some evidence that these include insects [75, 78]. However, this approach is the most complex of the three (cognition and computation-wise) as it requires the recognition of conspecifics combined with extrapolation capabilities.
- The second strategy (OMID) entirely ignores any partial information. This requires differentiating between fully vs. partially observed neighbors, which implies using recognition of conspecifics. However, it is somewhat simpler than COMPLID since it only filters out erroneous visual stimuli rather than computing the correct stimuli.
- The last strategy (PARTID) treats each segment of a neighbor as if it represents a full-length body. Hence, it is the simplest of the three since it requires minimal cognitive processing from the individual. It does not rely on dedicated conspecific recognition mechanism but instead clusters distinguishable visual features and regards each cluster as a neighbor, a relatively simple process.

However, the same simplicity also results in PARTID providing the most erroneous perception of the surrounding agent, as parts of neighbors’ segments change in their degree of visibility due to closer neighbors revealing or occluding them, which in turn is perceived as neighbors moving–quickly—away from or towards the focal agent (i.e., large absolute magnitude of the ***v***_***j***,***r***_ component).

The difference in the required computational power under the different approaches is very significant, as in nature, organisms demonstrating collective motion are very often limited in this respect (small brains, simple neuronal substrates). Hence, finding the least computationally demanding algorithm that is still capable of reaching flocking can potentially explain the actual mechanisms involved in the flocking of these relatively simple species. From this perspective, PARTID has the least requirements for visual information processing, while COMPLID has the most requirements.

In the torus arena, all three perception approaches of occlusions have successfully demonstrated the flocking transition from a disordered initial state to an ordered collective state. However, a detailed analysis reveals slight differences in convergence rates, where PARTID consistently appears to be slower to converge than the other strategies. This deficiency of PARTID is more pronounced at a higher density of neighbors, where occlusions are more frequent, and thus PARTID makes more errors.

When we evaluated the models in constrained environments (non-periodic bounds; corridor and ring), the general following conclusion emerges: *at best*, PARTID performs as well as others; *most often, it is far slower to increase order over time, and has lower long-term order parameter values*. Note that such constrained environments are common in nature. The topography of natural terrain has creeks, valleys, ridges, and other lateral constraints resulting in effectively constrained geometry. As is well established (see [44, 80] and references within), marching locust bands successfully maintain flock formation despite such constraints.

This raises a puzzle. On the one hand, an occlusion-handling method (PARTID) that is computationally cheap, and employs mechanisms whose existence in insects is generally accepted, is brittle and generally inferior to others exactly in the type of settings in which natural swarms, and in particular locust swarms, excel. On the other hand, in terms of order evolution over time, as well as order value at the end of the simulation, *COMPLID appears to be superior to the others in most cases in its convergence rate and long-term value and inferior to none*. However, COMPLID implies complex capabilities for recognizing conspecifics and for being able to extrapolate complete neighbor outlines from partial visual clues. While there is some limited evidence that insects are able to carry out such tasks (e.g., to extrapolating environment contours [75, 78]), recent laboratory studies of locust nymphs have demonstrated that they move in response to simulated movement of random visual patterns, which cannot be recognized as other locust [36].

Considering our results from constrained arenas, it is tempting to declare that PARTID is an oversimplification of the perceptive mechanisms in locust vision, and that advanced computational capabilities are required for coping with partially-occluded neighbors, as assumed by the proposed OMID or COMPLID approaches. However, examining related investigations offers other possibilities, as we discuss below.

### 5.2 Reliable Distance Estimation, Revisited

The critical weakness of all the models under the restricted perceptual capabilities we allow, is in the estimation of distance to neighbors. The geometry of the visual image denotes a single subtended angle *α*, parallel to the *horizontal* plane of motion (the plane on which the agent is moving) as the basis of distance estimation. As detailed in Section 2.1, we assume no information is given to the agent about the neigbor’s orientation. Lacking this information, the model assumes it is heading in a direction perpendicular to the LOS. Violations of this assumption insert errors into the distance estimations. Sans occlusions, their effects on the emerging flocking order is clear, but not dominant to the degree it prohibits flocking (consider the results in low-density arenas, for instance).

In the presence of partial occlusions, the errors caused by the assumptions of the model may gravely affect the result, *depending on the occlusion-handling strategy*. COMPLID relies on (assumed) complex capabilities of the agent to extrapolate the true dimensions of partially-occluded neighbors from visible parts. As a result, its distance estimates with respect to partially-occluded neighbors are the same as with fully-visible neighbors; it is therefore relatively robust to occlusions, in the sense that its performance should not change much as they become more frequent. In contrast, PARTID, which considers every visible part as an agent by itself, is gravely affected by partial occlusions. A small visible part of an occluded agent would be considered a distant neighbor. If the part grows—more of the occluded agent becomes visible, e.g., because the occluding agent in between is moving—then that same perceived distance agent now captures a much wider subtended angle, and would suddenly be perceived as close.

In other words, methods for *reliable* distance measurement in monocular images (other than those implied by COMPLOD and OMID) can help avoid the failures of PARTID. The complexity and biological plausibility of such methods should be considered vis-a-vis the processes assumed by strategies we already discuss above: neighbor contour extrapolation (COMPLID) and conspecifics recognition (COMPLID ahd OMID). Below, we discuss such potential methods, introduced in previous studies.

Several studies touch on the critical relationship between the agent morphology and distance estimation. Ignoring occlusions, visual perception of *circular* agents avoids the errors introduced by incorrect interpretation of *α*, as discussed in Section 2.1 and Appendix A.1. Moshtag et al. [58] and Berlinger et al. [85] demonstrated vision-based collective motion in physical robots, treating them as circles. Still, partial occlusions may cause rapid changes to *α*, and would make distance estimation unreliable under such conditions.

Bastien et al. [56] (and later Qi et al. [57]) demonstrated that under the assumption of circular agents, a completely different control approach can be taken, which avoids identifying individual neighbors or estimating the distance and heading of neighboring agents altogether. Rather, the agents only mark the projected blocking of the visual field by neighbors, without tracking them individually; angular segments in the field of view, blocked by neighbors, are marked as such, without a measurement of distance or identification of the neighbor. As a result, this approach is not sensitive to the occlusions in the same manner as the models introduced here.

While robots may be built to be circular in shape, natural swarming animals are most often elongated—with locusts being an example. As we were initially motivated by the behavior of natural swarms, the experiments above were tested using elongated simulated agents. Nonetheless, the reduction in errors offered by assuming a circular shape raises the question of the importance of the agent’s morphology to the presented models. To address this question, we experimented with different length-to-width ratios. The analysis (Appendix A.4) reveals that the performance of the different models varied *widely* when the body length ratio was changed, both in the rate by which order increases, as well as in long-term order values. Moreover, the qualitative relationships between models varied as well. In other words, the elongation of the agent has a dramatic effect on the emergence of ordered collective motion.

Consequently, there is a possible limitation to the models introduced in this study, as their behavior is dependent on the body length ratio, well as, undoubtedly, on other factors (visual range *R*, steering parameter *η*, etc.). Combining this understanding with the approach 678 taken by Bastien et al. [56] and Qi et al. [57] would be extremely interesting for future research, as it may enable robust collective motion for elongated shapes, avoiding the distance estimation errors caused by occlusions.

There are additional strategies that may be applicable. Because the agent’s shape is a given property in nature, how else might an agent overcome the errors introduced into its distance estimates by the variance in *α* (esp. with partial occlusions)?

First, we may attempt to infer the orientation of the agent, to improve the distance estimate, from enriched visual information. For instance, this may be done by matching distortions of known visual features to compute the orientation. Figure 13 illustrates an hypothetical example of how this might work with locust nymphs. Note that this type of process is still possible with flat 2D sensing (no height), as the distortions are revealed as distance changes between visual features of the known template.

**Fig 13.**
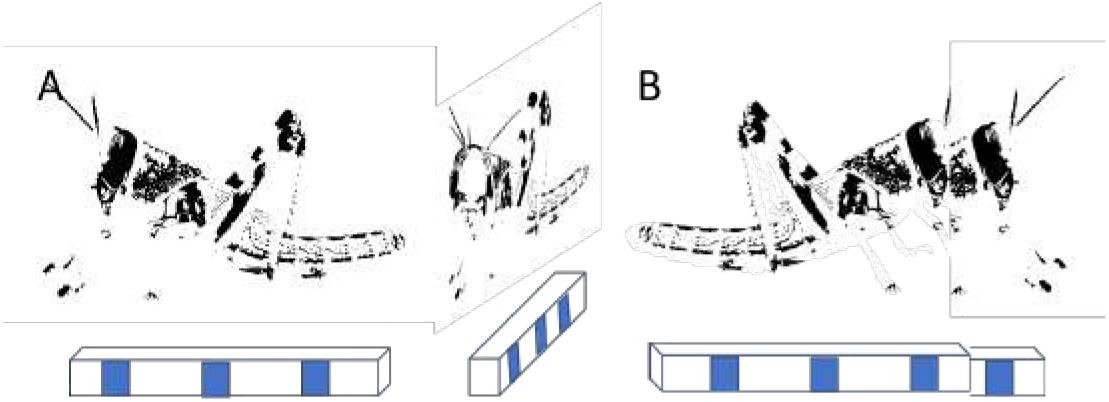
Hypothetical example of how recognized template distortions may be used to infer orientation of visible and partially-visible neighboring locust nymphs based on its black patterning alone (i.e., not fully-detailed conspecific recognition). **A**. Template distortions due to different headings. **B**. Template distortions due to partial occlusion.

Second, independently, we may remove the artificial restriction on perception of a flat world, and consider the more realistic view that the agent views three-dimensional objects. Visible neighbors would then be characterized by *two subtended angles*, one measuring the horizontal dimension of the neighbor (the familiar *α* subtended angle), and one measuring its vertical dimension, i.e., its height (let us call it *γ*). Note that for elongated agents moving on the horizontal plane, *α* depends heavily on the orientation of the observed agent, but *γ* does not. For example, in Figure 13-A, note how the subtended angle of the neighbor *α* changes with its heading, much more than its height *γ*. Integrating this information enables much more robust distance estimations, and as both natural agents and robots move in three-dimensional worlds, it is commonly applicable [54, 85]. The use of *γ* can alleviate the errors caused by partial occlusions considerably when neighbors’ height is visible while their horizontal dimension is partially hidden.

Third, we may attempt to generate depth information from monocular images taken over time. In computer science, this is called *structure from motion* (SfM), a complex process that generates depth information (and thus, estimated distance) from multiple images taken by a single moving camera, at different (close) times [86, 87]. While this is typically carried out in a static environment (i.e., the agent is localized with respect to static objects), it is possible in principle to apply this also to moving neighbors (see below for a discussion of the analogous challenge for optical flow generation).

None of the approaches discussed above for distance estimation from monocular images, completely solves the problem raised by partial occlusions. However, independently or in combination, they may alleviate it to an extent that enables computationally-simpler mechanisms to perform as well as those requiring complex processes. Indeed, more generally, allowing for rich visual projected information allows more robust measurements, based on many visual features, including shading, 3D shapes, color and spectral data, texture, etc. [31]. Even relatively simple combinations of visual features can be very useful. For example, Wang et al. [59] demonstrated implicit cooperation between robotic swarm members using visual inputs. Utilizing specific schemes for positioning poles holding specifically-placed sets of LED lights, the robots were able to estimate the relative positioning, velocity, state, and other features of their neighbors. Royer et al. [39] and Dong et al. [34] survey the progress in this direction in robotics. There has also been great interest recently in applying machine learning approaches to the challenge of estimating depth from monocular images, utilizing data containing rich visual information [37, 88]. These studies, rooted in robotics and engineering, may inspire investigations into biologically-plausible counterparts.

### Reliable Velocity Estimates

A common attractive component in all the models we presented is that of their reliance on optical flow as a key step in measuring 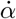 and 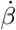. The optical flow is the vectorial difference between the same features or object in two images. It is generally accepted that optical flow is employed in nature by many species, including insects [33, 38]. It is certainly an important technique in robots [35, 89, 90].

One of the most common challenges to the use of optical flow in crowded environments, even ignoring the issue of occlusions, is that it is difficult to compute when the agent’s social environment is moving independently of a moving observer. In other words, distinguishing the optical flow of observed agents that are moving in the vicinity of the observer, while the observer itself is moving, is computationally difficult, prone to errors, and sometimes impossible (this challenge also arises for SfM processes, discussed above). As we conducted simulation experiments in which estimations were produced by a simulated process, we could ignore this complexity. Employing the reductionist model in robots—or investigating its potential use in nature—would require tackling this computation; neither animals nor robots can side-step this issue.

We note that computing optical flow when either the observer is moving (and others are standing still), or when the observer halts (and others are moving) is relatively easy. However, for the purposes of employing the model we present here, neither simplified variant would appear sufficient, as agents move while observing. In this context, it is important to note that previous work has established that the Pause-and-Go motion scheme plays a role in the repeated decision-making of locusts in a swarm [21, 26, 91], i.e., a representation of the local environment, utilized for deciding whether and in what direction to move, is constructed by the locusts when standing.

Assuming locusts do rely on optical flow in their collective motion decisions, we raise the hypothesis that locust reduce the complexity of the optical flow by carrying it out in two stages: pausing briefly, they compute the optical flow of their social environment to estimate the neighbor’s motion: then, moving briefly, they estimate their own velocity vector from the optical flow, assuming the world as static. We intend to examine this hypothesis in future investigations in locusts, as well as in robotic models.

The research presented above has studied different aspects of vision-based collective motion in swarms. The biological inspiration was to study visual, non-stereoscopic inputs, without direct distance measurements and while accounting for occlusions. Our primary quantitative “lens” for this investigation is the polarization measure of order (defined in Eq. 10), which is commonly and frequently used in studies of collective motion research [14, 20, 81–83]. Using this order measure, we have shown that the reductionist model is sufficient in many cases to achieve ordered collective behavior in a swarm. It is possible that perhaps some other types of measures of order could reveal additional information about these different cases.

The key insights we offer are that the behavior of a swarm can utilize a very simple, monocular visual perception to achieve robust flocking within the constraints of the agent’s body shape and wide field-of-view, and that the flocking process properties depend upon the particular mechanisms used to address occlusions. This is an important area of continued investigation, as it affects not only visual perception, but also other sensory modalities [28]. More generally, we hope that the study presented here contributes to the understanding of the fascinating interrelations between agent morphology, collective motion control algorithms, and visual perception capabilities. We also hope that it underscores questions requiring further investigations.

## Acknowledgments

We gratefully acknowledge partial funding support by ISF Grant #2306/18. We would like to thank Dr. Michael Krongauz for his invaluable contribution to this research and the development of the model. Thanks to K. Ushi.

## A Appendix: Supplementary materials

This appendix provides additional details and information accompanying, clarifying, and illustrating the main body of results reported in the study.

## A.1 Measurement of distance to neighbors in monocular, non-stereoscopic vision

In this appendix, we briefly describe the computation of distance from the focal agent to a neighboring agent. This is described for two different morphologies. Firstly, in Fig. 14, the computation of distance from the subtended angle is shown for idealized circular-shaped agents. It can be seen that for circles, there exists a one-to-one relationship, given by 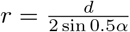 between the subtended angle and the distance. There is no ambiguity; hence the distance calculation is exact in the circular case.

**Fig 14.**
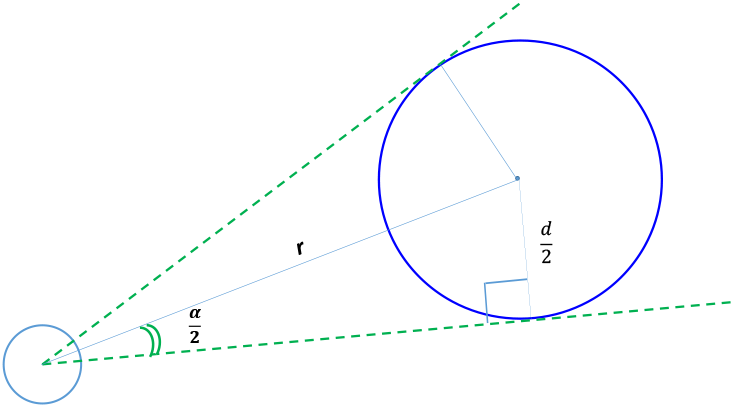
Exact distance r computation from subtended angle *α* for circular morphology. Assumption of a circular (or spherical in 3D) shape of the agents, enables precise computation of distance r to a neighbor while employing the non-stereoscopic visual parameter of subtended angle *α*. The small circle on the lower left depicts the focal agent’s visual sensor and the large circle on the right depicts a circular neighboring agent. Green lines represent the extreme rays toward the neighbor, as seen by the focal agent. The angle between the radius and the tangent extreme ray is always 90^0^ by geometrical definition. Therefore, as shown 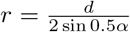 where d is the circle’s diameter and *α* the subtended angle measured.

**Fig 15.**
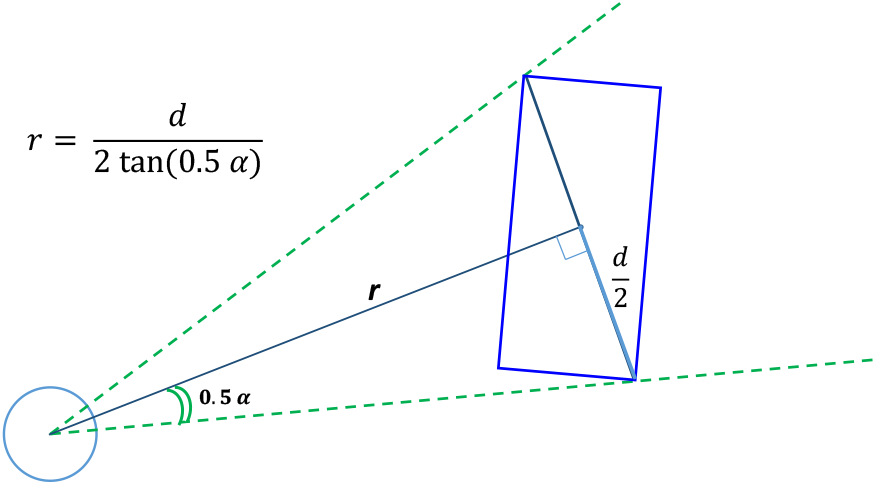
Approximate distance r computation from subtended angle *α* for elongated rectangles. Assumption of rectangular shape for neighboring agents, enables only an approximate computation of distance r to the neighbor, using non-stereoscopic visual parameter of the subtended angle *α*. The small circle on the lower left depicts the visual sensor of the focal agent, and the large circle on the right depicts a circular neighboring agent. Green lines are the edge rays towards the neighbor, as seen from the focal agent. The angle between the edge rays is the observed subtended angle *α*. Therefore as shown in the figure 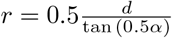

### A.2 Derivation of Equation (7)

We know (from Eq. (4)) that

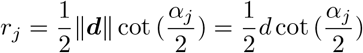

by differentiating Eq. (4) with respect to time *t*, we see (Eq. 6) that

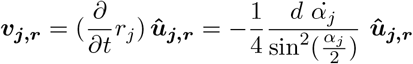

where 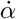 denotes the time derivative of the subtended angle.

Expressing 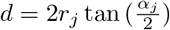 from Eq. (4) and substituting *d* into Eq. (6) results in the following derivation of the radial velocity ***v***_***j***,***r***_,

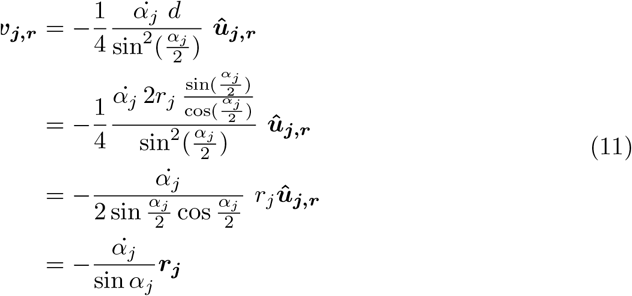

which is what appears in Eq. (7).

### A.3 Population Size *N* in the torus environment

We experimentally set the value of *N* in different arenas, to values that proved informative in the sense that they highlighted and clarified the differences between different strategies or other parameters. Below, we show the procedure and results used to set the values of *N* used in the torus arena. For other arenas, a similar experimental analysis was carried out.

We begin with the values that have already been set for the rest of the parameters: *R* = 3[BL], *η* = 0.01, length-to-width ratio set at 3. We then vary *N* . Figures 16a–16b, show the results of 50 independent trials. For relatively small *N* sizes, convergence to an ordered state is slow (at best) and probably non-existent. In the context of range-limited vision (*R* = 3[BL]), the sparse density undoubtedly inhibits overall convergence to ordered flocking. Increasing *N* leads to a higher long-term order parameter (Figure 16b), though clearly the rate of convergence differs (Figure 16a). Based on these results, we typically use population sizes *N* = 60, 120, 180 in the experiments in the torus arena.

**Fig 16.**
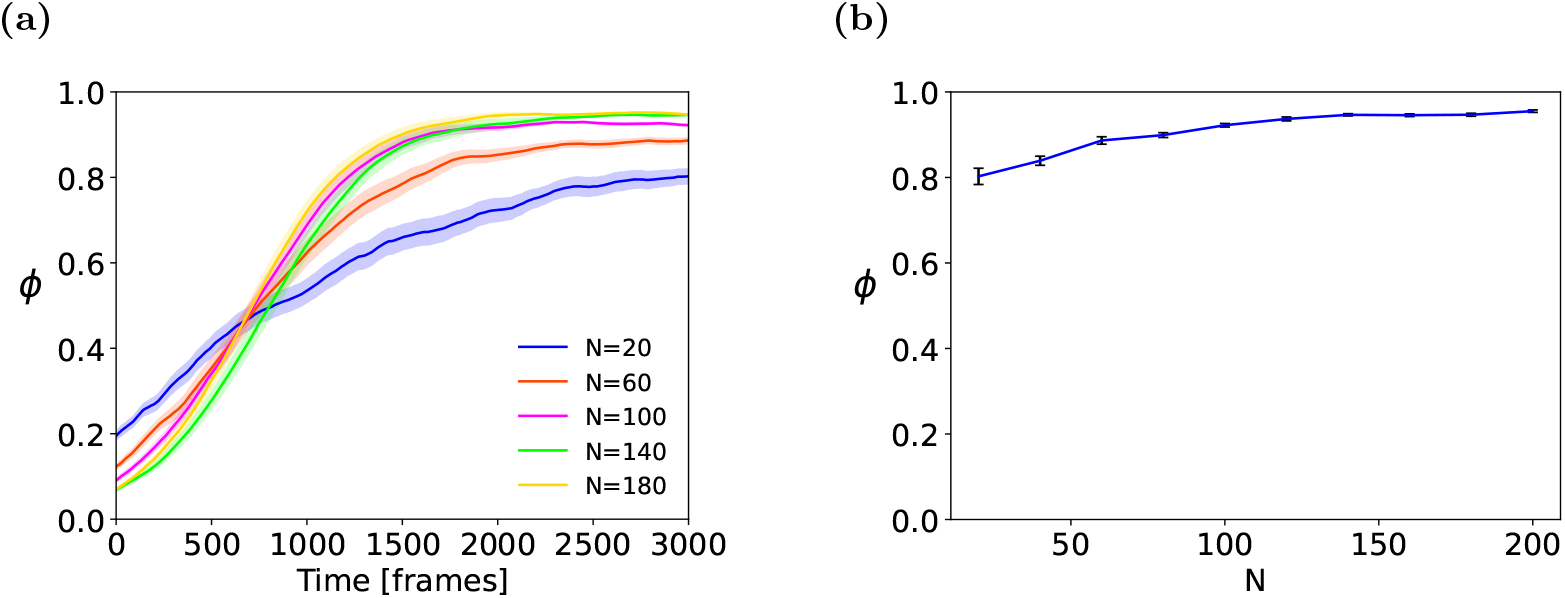
**(a)** Time-dependent and **(b)** long-term sensitivity analysis for visual range *N*, in the torus arena. Means and standard errors are shown for 50 trials.

### A.4 Influence of the Length-to-Width Ratio

We examine the influence of elongation on the emergence of order in flocking. In nature, nymph lengths vary as they grow to adulthood. A question also arises (when discussing the visual perception of the locust) regarding the inclusion of out-stretching legs in the perceived image. Based on our own measurements of the locust in our laboratory, we have chosen a body length-to-width ratio of 3 (i.e., length is 3 times the body width) as a baseline, which was used in the experiments reported above.

However, in response to an excellent comment from an anonymous reviewer of the study, we sought to examine whether these settings influence the results. In particular, different models exist for vision-based flocking, for the case where agents are circular [51, 56, 58]. If the models introduced in this study is insensitive to the length-to-width ratio, then perhaps these other models could be just as useful in informing our understanding of how flocking may work in locusts (or other species that are elongated).

We therefore experimented with other ratios: a ratio 1:1 (agents are perfect square), the 3:1 ratio, and a ratio of 6:1. Figures 17 and 18 report on the results from these experiments, in all environments (with the other parameters set as before: *N* = 100, *R* = 3, *η* = 0.01, etc.). In all, we tested the principal model, as well as all three occlusion-handling strategies. As before, we conducted 50 independent trials in each settings and present the means and standard errors. Overall, 50 ×4× 9 = 1800 simulation runs were executed in this experiment.

**Fig 17.**
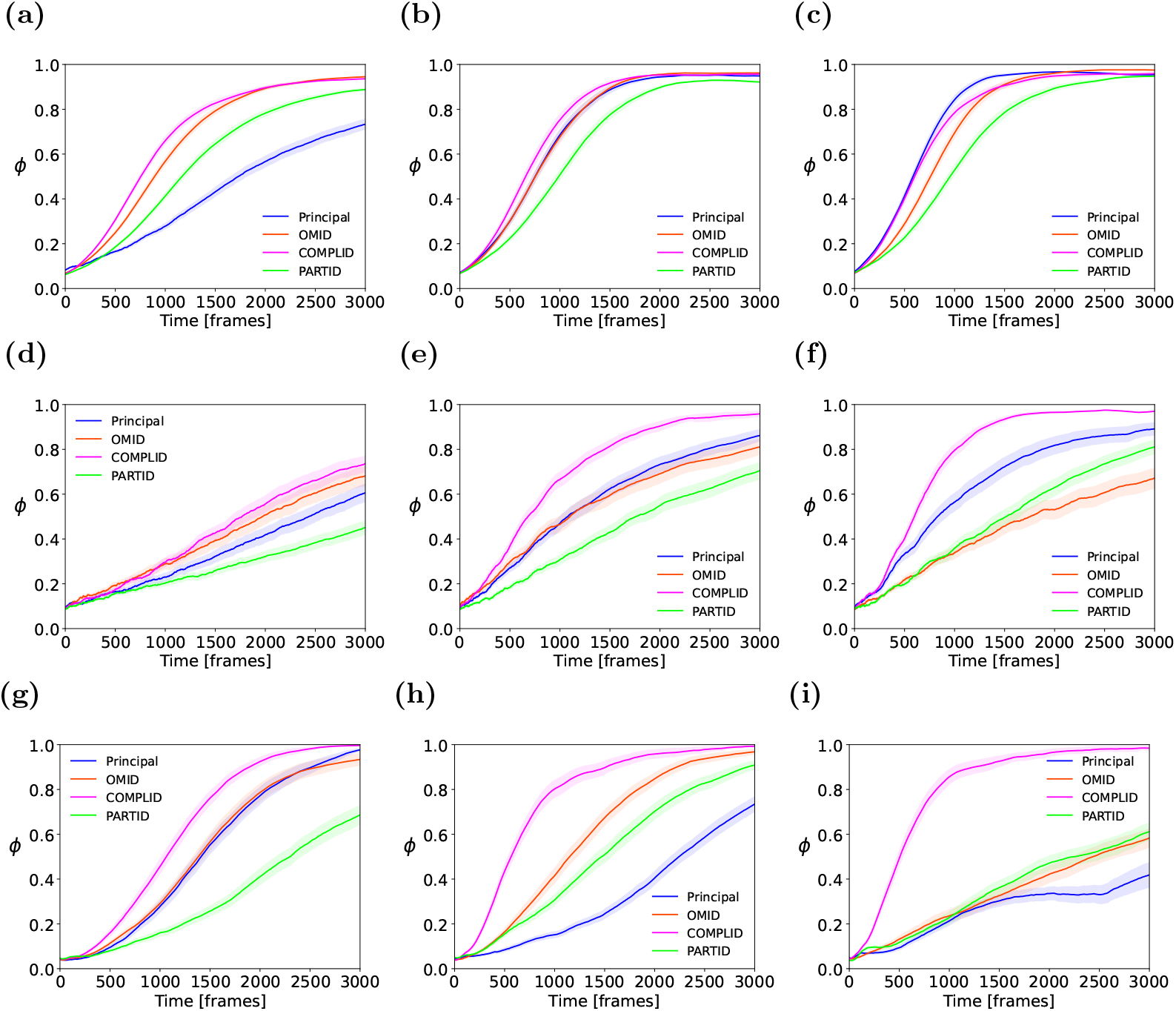
Order measure *ϕ* evolving in time *t* = 1 … 3000, for different body length-to-width ratios (left to right columns: 1:1, 3:1, 6:1) and arenas (top to bottom rows: torus, corridor, ring). The results are shown for the principal model, and the three occlusion-handling strategies. It is evident that the trend of order parameter evolution, for all models, depends significantly upon the length-to-width ratio.

**Fig 18.**
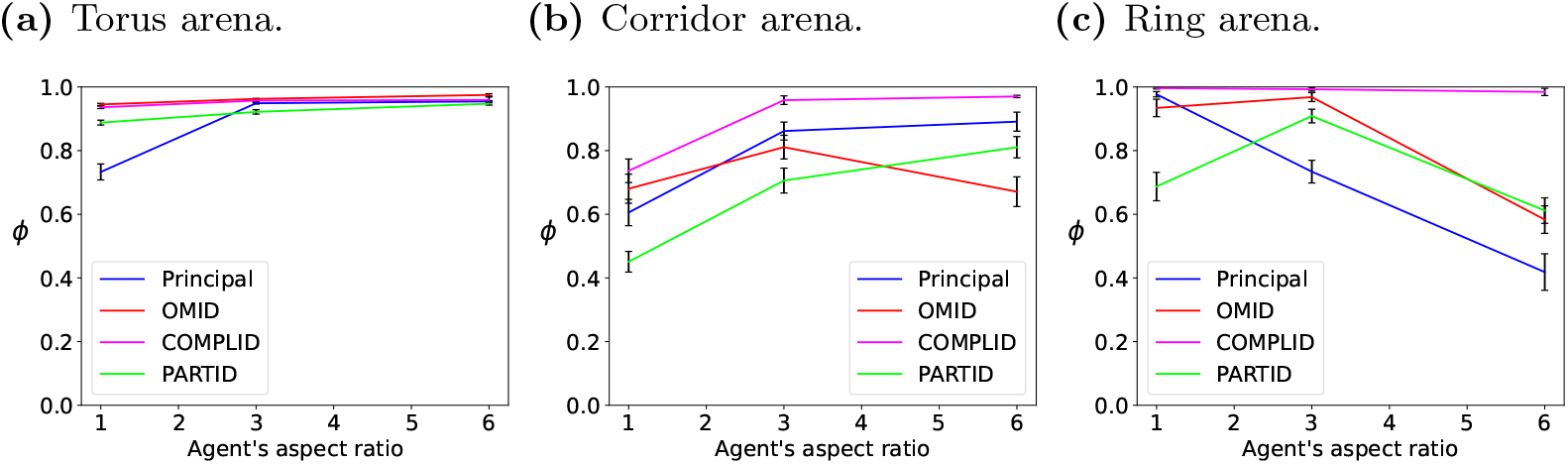
Long term order (*ϕ* at *t* = 3000), for different body length-to-width ratios (horizontal axis in each subfigure), for different arenas (see captions). The results are shown for the principal model, and the three occlusion-handling strategies. Mean values and standard errors shown for 50 independent trials.

Figure 17 shows the order parameter evolving over time in various settings. The subfigures are arranged by columns (different ratios) and rows (different arenas). The left column of the figures (Figures (a), (d), (g)) shows the results for ratio 1:1, the middle column (Figures (b), (e), (h)) show the results for the baseline ratio 3:1 used in the main group of experiments as reported above, and the rightmost column of figures (Figures (c), (f), (i)) show the results for ratio 6:1. The top row shows the results from the toroidal arena, the middle row shows results for the narrow corridor arena, and the bottom row shows the results for the narrow ring arena.

The figures show that the evolution of order, over time, is greatly influenced by the body length-to-width ratio. This is generally true for all the models, and thus we conclude that the body length ratio is an important factor in the rate of convergence. Qualitatively, we note that higher ratios (elongated morphology) improve the rate of order increase in both the torus and corridor arenas. We believe these less-constrained arenas are closer to natural environments than the small ring-shaped arena (where greater elongation reduces—-possibly eliminates-—the the rate of order increase over time.

Figure 18 presents the long-term order (*ϕ*, at time *t* = 3000) in the different arenas, for the different ratios; essentially it displays a cross-section at time *t* = 3000 of Figure 17. The results show that in general, application of the models in the torus arena is more robust with respect to body length-to-width ratios, but this does not hold true for the other arenas. Overall, we note that while the COMPLID model tends to outperform the others across arenas and length ratios, but this is a qualitative observation, and subject to future investigation. Here, just as in Figure 17, we reach the conclusion that any evaluation of the models in the context of a target species requires taking the body length-to-width ratio into account.

## Code

The source code for the model implementation in simulation is available at https://github.com/kronga/vision-based-collective-model.

## References

1. Ballerini M, Cabibbo N, Candelier R, Cavagna A, Cisbani E, Giardina I, et al. Interaction ruling animal collective behavior depends on topological rather than metric distance: Evidence from a field study. Proceedings of the National Academy of Sciences of the United States of America. 2008;105(4):1232–1237. doi:10.1073/PNAS.0711437105.

2. Ballerini M, Cabibbo N, Candelier R, Cavagna A, Cisbani E, Giardina I, et al. Empirical investigation of starling flocks: a benchmark study in collective animal behaviour. Animal Behaviour. 2008;76(1):201–215. doi:10.1016/J.ANBEHAV.2008.02.004.

3. Rosenthal SB, Twomey CR, Hartnett AT, Wu HS, Couzin ID. Revealing the hidden networks of interaction in mobile animal groups allows prediction of complex behavioral contagion. Proceedings of the National Academy of Sciences of the United States of America. 2015;112(15):4690–4695. doi:10.1073/PNAS.1420068112.

4. Handegard NO, Boswell KM, Ioannou CC, Leblanc SP, Tjostheim DB, Couzin ID. The Dynamics of Coordinated Group Hunting and Collective Information Transfer among Schooling Prey. Current Biology. 2012;22(13):1213–1217. doi:10.1016/J.CUB.2012.04.050.

5. Buhl J, Sumpter DJT, Couzin ID, Hale JJ, Despland E, Miller ER, et al. From disorder to order in marching locusts. Science. 2006;312(5778):1402–1406. doi:10.1126/SCIENCE.1125142/SUPPLFILE/BUHL.SOM.PDF.

6. Uvarov B, et al. Grasshoppers and locusts. A handbook of general acridology Vol. 2. Behaviour, ecology, biogeography, population dynamics. Centre for Overseas Pest Research; 1977.

7. Zhang HP, Be’er A, Florin EL, Swinney HL. Collective motion and density fluctuations in bacterial colonies. Proceedings of the National Academy of Sciences of the United States of America. 2010;107(31):13626–13630. doi:10.1073/PNAS.1001651107.

8. Henderson LF. The statistics of crowd fluids. Nature. 1971;229:381–383.

9. Wolff M. Notes on the behaviour of pedestrians. In: Birenbaum A, Sagarin E, editors. People in Places: The Sociology of the Familiar. Nelson; 1973. p. 35–48,.

10. Helbing D, Molnar P, Farkas IJ, Bolay K. Self-organizing pedestrian movement. Environment and Planning B. 2001;28:361–384.

11. Daamen W, Hoogendoorn SP. Experimental research of pedestrian walking behavior. Transportation Research Record. 2003; p. 20–30,.

12. Kaminka GA, Fridman N. Simulating Urban Pedestrian Crowds of Different Cultures. ACM Transactions on Intelligent Systems and Technology. 2018;9(3):27:1–27:27. doi:10.1145/3102302.

13. Reynolds CW. Flocks, herds and schools: A distributed behavioral model. In: Proceedings of the 14th annual conference on Computer graphics and interactive techniques; 1987. p. 25–34.

14. Vicsek T, Czirk A, Ben-Jacob E, Cohen I, Shochet O. Novel Type of Phase Transition in a System of Self-Driven Particles. Physical Review Letters. 1995;75(6):1226. doi:10.1103/PhysRevLett.75.1226.

15. Crowd simulation software; 2004.

16. Fridman N, Kaminka GA. Modeling Pedestrian Crowd Behavior Based on a Cognitive Model of Social Comparison Theory. Computational and Mathematical Organizational Theory. 2010;16(4):348–372.

17. Tsai J, Fridman N, Brown M, Ogden A, Rika I, Wang X, et al. ESCAPES -Evacuation Simulation with Children, Authorities, Parents, Emotions, and Social comparison. In: Proceedings of the Tenth International Joint Conference on Autonomous Agents and Multi-Agent Systems (AAMAS-11); 2011.

18. Hamann H. Swarm Robotics: A Formal Approach. Springer; 2018.

19. Deutsch A, Theraulaz G, Vicsek T. Collective motion in biological systems; 2012.

20. Vicsek T, Zafeiris A. Collective motion; 2012.

21. Ariel G, Ayali A. Locust Collective Motion and Its Modeling. PLOS Computational Biology. 2015;11(12):e1004522. doi:10.1371/JOURNAL.PCBI.1004522.

22. Kolpas A, Moehlis J, Kevrekidis IG. Coarse-grained analysis of stochasticity-induced switching between collective motion states. Proceedings of the National Academy of Sciences of the United States of America. 2007;104(14):5931–5935. doi:10.1073/PNAS.0608270104.

23. Escudero C, Yates CA, Buhl J, Couzin ID, Erban R, Kevrekidis IG, et al. Ergodic directional switching in mobile insect groups. Physical Review E -Statistical, Nonlinear, and Soft Matter Physics. 2010;82(1):011926. doi:10.1103/PHYSREVE.82.011926/FIGURES/1/MEDIUM.

24. Aoki I. internal Dynamics of Fish Schools in Relation to Inter-fish Distance. Nippon Suisan Gakkaishi. 1984;50(5):751–758. doi:10.2331/suisan.50.751.

25. Bode NWF, Faria JJ, Franks DW, Krause J, Wood AJ. How perceived threat increases synchronization in collectively moving animal groups. Proceedings of the Royal Society B: Biological Sciences. 2010;277(1697):3065–3070. doi:10.1098/RSPB.2010.0855.

26. Ariel G, Ophir Y, Levi S, Ben-Jacob E, Ayali A. Individual Pause-and-Go Motion Is Instrumental to the Formation and Maintenance of Swarms of Marching Locust Nymphs. PLOS ONE. 2014;9(7):e101636. doi:10.1371/JOURNAL.PONE.0101636.

27. Cucker F, Smale S. Emergent behavior in flocks. IEEE Transactions on automatic control. 2007;52(5):852–862.

28. Kunz H, Hemelrijk CK. Simulations of the social organization of large schools of fish whose perception is obstructed. Applied Animal Behaviour Science. 2012;138(3-4):142–151. doi:10.1016/J.APPLANIM.2012.02.002.

29. Mascalzoni E, Regolin L. Animal visual perception. Wiley Interdisciplinary Reviews: Cognitive Science. 2011;2(1):106–116. doi:10.1002/WCS.97.

30. Goldstein EB. Encyclopedia of perception. Sage; 2009.

31. Ma Y, Soatto S, Košecká J, Sastry S. An invitation to 3-D vision: from images to geometric models. vol. 26. Springer; 2004.

32. Nityananda V, Read JC. Stereopsis in animals: evolution, function and mechanisms. Journal of Experimental Biology. 2017;220(14):2502–2512.

33. Hamada T. Vision, action, and navigation in animals. Visual Navigation: From Biological Systems to Unmanned Ground Vehicles. 1997;2:1.

34. Dong X, Garratt MA, Anavatti SG, Abbass HA. Towards real-time monocular depth estimation for robotics: A survey. IEEE Transactions on Intelligent Transportation Systems. 2022;23(10):16940–16961.

35. Serres JR, Ruffier F. Optic flow-based collision-free strategies: From insects to robots. Arthropod structure & development. 2017;46(5):703–717.

36. Bleichman I, Yadav P, Ayali A. Visual processing and collective motion-related decision-making in desert locusts. Proceedings of the Royal Society B. 2023;290(1991):20221862.

37. Ming Y, Meng X, Fan C, Yu H. Deep learning for monocular depth estimation: A review. Neurocomputing. 2021;438:14–33.

38. Srinivasan M, Zhang S, Lehrer M. Honeybee navigation: odometry with monocular input. Animal behaviour. 1998;56(5):1245–1259.

39. Royer E, Lhuillier M, Dhome M, Lavest JM. Monocular vision for mobile robot localization and autonomous navigation. International Journal of Computer Vision. 2007;74(3):237–260.

40. Egelhaaf M, Kern R. Vision in flying insects. Current opinion in neurobiology. 2002;12(6):699–706.

41. Ayali A. The puzzle of locust density-dependent phase polyphenism. Current opinion in insect science. 2019;35:41–47.

42. Cullen DA, Cease AJ, Latchininsky AV, Ayali A, Berry K, Buhl J, et al. From molecules to management: mechanisms and consequences of locust phase polyphenism. In: Advances in Insect Physiology. vol. 53. Elsevier; 2017. p. 167–285.

43. Zhang L, Lecoq M, Latchininsky A, Hunter D. Locust and grasshopper management. Annu Rev Entomol. 2019;64(1):15–34.

44. Dkhili J, Berger U, Hassani LMI, Ghaout S, Peters R, Piou C. Self-organized spatial structures of locust groups emerging from local interaction. Ecological Modelling. 2017;361:26–40.

45. Bazazi S, Buhl J, Hale JJ, Anstey ML, Sword GA, Simpson SJ, et al. Collective motion and cannibalism in locust migratory bands. Current biology. 2008;18(10):735–739.

46. Knebel D, Ayali A, Guershon M, Ariel G. Intra-versus intergroup variance in collective behavior. Science advances. 2019;5(1):eaav0695.

47. Pita D, Collignon B, Halloy J, Fernández-Juricic E. Collective behaviour in vertebrates: A sensory perspective. Royal Society Open Science. 2016;3(11). doi:10.1098/RSOS.160377.

48. Lemasson BH, Anderson JJ, Goodwin RA. Collective motion in animal groups from a neurobiological perspective: The adaptive benefits of dynamic sensory loads and selective attention. Journal of Theoretical Biology. 2009;261(4):501–510. doi:10.1016/J.JTBI.2009.08.013.

49. Strandburg-Peshkin A, Twomey CR, Bode NWF, Kao AB, Katz Y, Ioannou CC, et al. Visual sensory networks and effective information transfer in animal groups. Current Biology. 2013;23(17):R709–R711. doi:10.1016/J.CUB.2013.07.059/ATTACHMENT/9C177EE8-47CA-4A66-BCD2-FBB3BDE6028B/MMC3.MP4.

50. Lemasson BH, Anderson JJ, Goodwin RA. Motion-guided attention promotes adaptive communications during social navigation. Proceedings of the Royal Society B: Biological Sciences. 2013;280(1754). doi:10.1098/RSPB.2012.2003.

51. Schilling F, Soria E, Floreano D. On the Scalability of Vision-Based Drone Swarms in the Presence of Occlusions. IEEE Access. 2022;10:28133–28146. doi:10.1109/ACCESS.2022.3158758.

52. Kaminka GA, Schechter-Glick R, Sadov V. Using Sensor Morphology for Multi-Robot Formations. IEEE Transactions on Robotics. 2008; p. 271–282.

53. Kaminka GA, Lupu I, Agmon N. Construction of Optimal Control Graphs in Multi-Robot Systems. In: Berman S, Gauci M, Frazzoli E, Kolling A, Gross R, Martinoli A, et al., editors. 13th International Symposium on Distributed Autonomous Robotic Systems (DARS-2016). Springer; 2016.

54. Collignon B, Séguret A, Halloy J. A stochastic vision-based model inspired by zebrafish collective behaviour in heterogeneous environments. Royal Society Open Science. 2015;3(1). doi:10.1098/RSOS.150473.

55. Soria E, Schiano F, Floreano D. The Influence of Limited Visual Sensing on the Reynolds Flocking Algorithm. In: Proceedings of the 3rd IEEE International Conference on Robotic Computing (IRC).Institute of Electrical and Electronics Engineers Inc.; 2019. p. 138–145.

56. Bastien R, Romanczuk P. A model of collective behavior based purely on vision. Science Advances. 2020;6(6). doi:10.1126/SCIADV.AAY0792/SUPPLFILE/AAY0792SM.PDF.

57. Qi J, Bai L, Xiao Y, Wei Y, Wu W. The emergence of collective obstacle avoidance based on a visual perception mechanism. Information Sciences. 2022;582:850–864. doi:10.1016/J.INS.2021.10.039.

58. Moshtagh N, Michael N, Jadbabaie A, Daniilidis K. Vision-based, distributed control laws for motion coordination of nonholonomic robots. IEEE Transactions on Robotics. 2009;25(4):851–860. doi:10.1109/TRO.2009.2022439.

59. Wang X, Wang F, Nie Z, Ai Y, Hu T. optiSwarm: Optical Swarm Robots using Implicit Cooperation. IEEE Sensors Journal. 2022;.

60. Judge SJ, Rind FC. The locust DCMD, a movement-detecting neurone tightly tuned to collision trajectories. Journal of Experimental Biology. 1997;200(16):2209–2216. doi:10.1242/JEB.200.16.2209.

61. Gray JR, Blincow E, Robertson RM. A pair of motion-sensitive neurons in the locust encode approaches of a looming object. Journal of Comparative Physiology A: Neuroethology, Sensory, Neural, and Behavioral Physiology. 2010;196(12):927–938. doi:10.1007/S00359-010-0576-7/FIGURES/8.

62. Bass M. Handbook of optics: volume i-geometrical and physical optics, polarized light, components and instruments. McGraw-Hill Education; 2010.

63. Goodale MA, Ellard CG, Booth L. The role of image size and retinal motion in the computation of absolute distance by the Mongolian gerbil (Meriones unguiculatus). Vision research. 1990;30(3):399–413. doi:10.1016/0042-6989(90)90082-V.

64. Santer RD, Rind FC, Stafford R, Simmons PJ. Role of an identified looming-sensitive neuron in triggering a flying locust’s escape. Journal of Neurophysiology. 2006;95(6):3391–3400. doi:10.1152/JN.00024.2006.

65. Ben-Nun A, Ayali A. Self body-size perception in an insect. Naturwissenschaften. 2013;100(5):479–484.

66. De Vries SEJ, Clandinin TR. Loom-sensitive neurons link computation to action in the Drosophila visual system. Current Biology. 2012;22(5):353–362. doi:10.1016/J.CUB.2012.01.007.

67. Bennett LV, Symmons PM. A review of estimates of numbers in some types of desert locust (Schistocerca gregaria (Forsk.)) populations. Bulletin of Entomological Research. 1972;61(4):637–649.

68. Luck SJ, Ford MA. On the role of selective attention in visual perception. Proceedings of the National Academy of Sciences. 1998;95(3):825–830.

69. Canosa RL. Real-world vision: Selective perception and task. ACM Transactions on Applied Perception (TAP). 2009;6(2):1–34.

70. Dunbier JR, Wiederman SD, Shoemaker PA, O’Carroll DC. Facilitation of dragonfly target-detecting neurons by slow moving features on continuous paths. Frontiers in Neural Circuits. 2012;0(OCTOBER 2012):1–29. doi:10.3389/FNCIR.2012.00079/BIBTEX.

71. Wang H, Peng J, Yue S. A Directionally Selective Small Target Motion Detecting Visual Neural Network in Cluttered Backgrounds. IEEE Transactions on Cybernetics. 2020;50(4):1541–1555. doi:10.1109/TCYB.2018.2869384.

72. Kanizsa G, Renzi P, Conte S, Compostela C, Guerani L. Amodal completion in mouse vision. Perception. 1993;22(6):713–721. doi:10.1068/p220713.

73. Singh M. Modal and amodal completion generate different shapes. Psychological Science. 2004;15(7):454–459. doi:10.1111/j.0956-7976.2004.00701.x.

74. Bruce V, Green PR, Georgeson MA. Visual perception: Physiology, psychology, & ecology. 4th ed. Hove & London: Psychology Press; 2003.

75. Nieder A. Seeing more than meets the eye: processing of illusory contours in animals. Journal of Comparative Physiology A 2002 188:4. 2002;188(4):249–260. doi:10.1007/S00359-002-0306-X.

76. Lin IR, Chiao CC. Visual equivalence and amodal completion in cuttlefish. Frontiers in Physiology. 2017;8(FEB):40. doi:10.3389/FPHYS.2017.00040/BIBTEX.

77. Cox MA, Schmid MC, Peters AJ, Saunders RC, Leopold DA, Maier A. Receptive field focus of visual area V4 neurons determines responses to illusory surfaces. Proceedings of the National Academy of Sciences of the United States of America. 2013;110(42):17095–17100. doi:10.1073/PNAS.1310806110.

78. Horridge GA, Zhang S, O’Carroll D. Insect perception of illusory contours. Philosophical Transactions of the Royal Society of London Series B: Biological Sciences. 1992;337(1279):59–64.

79. Schmidt E. Ernst Schmidt -Coding;. Available from: http://www.ernst-schmidt.com.

80. Amichay G, Ariel G, Ayali A. The effect of changing topography on the coordinated marching of locust nymphs. PeerJ. 2016;4:e2742.

81. Czirók A, Barabási AL, Vicsek T. Collective Motion of Self-Propelled Particles: Kinetic Phase Transition in One Dimension. Physical Review Letters. 1999;82(1):209. doi:10.1103/PhysRevLett.82.209.

82. Topaz CM, Ziegelmeier L, Halverson T. Topological data analysis of biological aggregation models. PloS one. 2015;10(5):e0126383.

83. Wang B, Wu Y, Wang G, Liu L, Chen G, Zhang HT. Transition in collective motion decision making. Phys Rev E. 2022;106:014611. doi:10.1103/PhysRevE.106.014611.

84. Keidar M, Kaminka GA. Efficient Frontier Detection for Robot Exploration. IJRR. 2014;33(2):215–236.

85. Berlinger F, Gauci M, Nagpal R. Implicit coordination for 3D underwater collective behaviors in a fish-inspired robot swarm. Science Robotics. 2021;6(50). doi:10.1126/SCIROBOTICS.ABD8668/SUPPLFILE/ABD8668SM.PDF.

86. Ullman S. The Interpretation of Structure from Motion. Massachusetts Institute of Technology; 1976. 476.

87. Ozyesil O, Voroninski V, Basri R, Singer A. A Survey of Structure from Motion; 2017.

88. Zhao C, Sun Q, Zhang C, Tang Y, Qian F. Monocular depth estimation based on deep learning: An overview. Science China Technological Sciences. 2020;63(9):1612–1627.

89. Sobey P. Active navigation with a monocular robot. Biological Cybernetics. 1994;71(5):433–440.

90. Zhan Q, Huang S, Wu J. Automatic navigation for a mobile robot with monocular vision. In: 2008 IEEE Conference on Robotics, Automation and Mechatronics. IEEE; 2008. p. 1005–1010.

91. Knebel D, Sha-ked C, Agmon N, Ariel G, Ayali A. Collective motion as a distinct behavioral state of the individual. iScience. 2021;24(4):102299. doi:10.1016/J.ISCI.2021.102299.

